# Yeast eIF2A has a minimal role in translation initiation and uORF-mediated translational control *in vivo*

**DOI:** 10.1101/2023.10.06.561292

**Authors:** Swati Gaikwad, Fardin Ghobakhlou, Hongen Zhang, Alan G. Hinnebusch

## Abstract

Initiating translation of most eukaryotic mRNAs depends on recruitment of methionyl initiator tRNA (Met-tRNAi) in a ternary complex (TC) with GTP-bound eukaryotic initiation factor 2 (eIF2) to the small (40S) ribosomal subunit, forming a 43S preinitiation complex (PIC) that attaches to the mRNA and scans the 5’-untranslated region (5’ UTR) for an AUG start codon. Previous studies have implicated mammalian eIF2A in GTP-independent binding of Met-tRNAi to the 40S subunit and its recruitment to specialized mRNAs that do not require scanning, and in initiation at non-AUG start codons, when eIF2 function is attenuated by phosphorylation of its α-subunit during stress. The role of eIF2A in translation *in vivo* is poorly understood however, and it was unknown whether the conserved ortholog in budding yeast can functionally substitute for eIF2. We performed ribosome profiling of a yeast deletion mutant lacking eIF2A and isogenic wild-type (WT) cells in the presence or absence of eIF2α phosphorylation induced by starvation for amino acids isoleucine and valine. Whereas starvation of WT confers changes in translational efficiencies (TEs) of hundreds of mRNAs, the *eIF2AΔ* mutation conferred no significant TE reductions for any mRNAs in non-starved cells, and it reduced the TEs of only a small number of transcripts in starved cells containing phosphorylated eIF2α. We found no evidence that eliminating eIF2A altered the translation of mRNAs containing putative IRES elements, or harboring uORFs initiated by AUG or near-cognate start codons, in non-starved or starved cells. Thus, very few mRNAs (possibly only one) appear to employ eIF2A for Met-tRNAi recruitment in yeast cells, even when eIF2 function is attenuated by stress.

## INTRODUCTION

Eukaryotic mRNAs are generally translated by the scanning mechanism, which commences with assembly of a 43S preinitiation complex (PIC) containing the small (40S) ribosomal subunit, a ternary complex (TC) of GTP-bound eukaryotic initiation factor 2 (eIF2) and methionyl initiator tRNA (Met-tRNAi), along with various other initiation factors. The 43S PIC attaches to the 5′-end of the mRNA, activated by eIF4F bound to the m^7^G cap (comprised of cap-binding protein eIF4E, scaffolding subunit eIF4G, and DEAD-box RNA helicase eIF4A) and scans the 5’ UTR to identify the AUG start codon, using complementarity with the anticodon of Met-tRNAi to recognize the AUG triplet. The 48S PIC arrested at the start codon joins with the large (60S) subunit to form an 80S initiation complex ready to synthesize the first peptide bond in protein synthesis (reviewed in (Hinnebusch 2014; Shirokikh and Preiss 2018)). A key mechanism for down-regulating bulk translation initiation during stress entails phosphorylation of the α-subunit of eIF2, which converts eIF2-GDP from substrate to inhibitor of its guanine nucleotide exchange factor, eIF2B, diminishing TC assembly. Translation of specialized mRNAs encoding certain stress-activated transcription factors regulated by inhibitory upstream open reading frames (uORFs) in their 5’-UTRs is induced rather than inhibited by eIF2α phosphorylation because the reduction in TC levels allows scanning PICs to bypass the uORF start codons and initiate further downstream at the start codon for the transcription factor coding sequences (CDS). This mechanism, dubbed the Integrated Stress Response in mammals, governs the translational induction of *GCN4* mRNA in budding yeast in response to amino acid starvation, dependent on the sole yeast eIF2α kinase, Gcn2. The Gcn4 protein thus induced activates the transcription of multiple genes encoding amino acid biosynthetic enzymes (reviewed in (Hinnebusch 2005; Gunisova et al. 2018; Dever et al. 2023)).

Various studies using cultured mammalian cells have implicated the auxiliary initiation factor eIF2A in translation initiation under stress conditions where eIF2 function is reduced by phosphorylation, in non-canonical initiation events where an internal ribosome entry site (IRES) bypasses the scanning mechanism for selecting AUG codons, or when a near-cognate codon (NCC) rather than AUG serves as initiation site (reviewed in (Komar and Merrick 2020)). eIF2A was originally characterized as a factor purified from rabbit reticulocytes that could stimulate GTP-independent recruitment of Met-tRNAi to the 40S subunit in response to an AUG codon; but could not stimulate the same reaction on native globin mRNA (Adams et al. 1975). Later studies suggested that the stimulation of Met-tRNAi binding was actually conferred by a protein co-purifying with eIF2A, identified as ligatin/eIF2D (Dmitriev et al. 2010; Skabkin et al. 2010), which was subsequently implicated in ribosome recycling at termination codons (reviewed in (Dever and Green 2012)). Experiments in cultured cells suggested that eIF2A could functionally substitute for eIF2 in recruiting Met-tRNAi to the AUG start codons of viral mRNAs, including Sindbis virus 26S mRNA (Ventoso et al. 2006) and HCV mRNA (Kim et al. 2011), which is driven by an IRES, when eIF2α is phosphorylated in the virus-infected cells. Other evidence suggested that eIF2A functionally cooperates with eIF5B in recruiting Met-tRNAi to the 40S-HCV IRES preinitiation complex (Kim et al. 2018); and it was implicated in IRES-dependent translation of c-SRC mRNA (Kwon et al. 2017). However, several independent studies did not support a role for eIF2A in initiation on Sindbis or HCV mRNAs (Jaafar et al. 2016; Sanz et al. 2017; Gonzalez-Almela et al. 2018).

Other lines of evidence implicated mammalian eIF2A in initiation at CUG initiation codons decoded by a leucyl tRNA versus Met-tRNAi in producing polypeptides presented by major histocompatibility complex (MHC) class I molecules (Starck et al. 2012); and in initiation at an upstream CUG codon to produce an N-terminally extended form of PTENα (Liang et al. 2014). eIF2A was also reported to support non-AUG initiation events involved in translation through an expansion of GGGGGC repeats present in the C9ORF72 gene in neuronal cells exhibiting elevated eIF2α phosphorylation (Sonobe et al. 2018). There is additional evidence that eIF2A mediates initiation at UUG- or CUG-initiated uORFs in the 5’UTR of BiP mRNA that appear to enhance BiP translation when eIF2α phosphorylation is induced by ER stress (Starck et al. 2016). eIF2A has also been implicated in stimulating translation of positive-acting uORFs initiated by CUG or other NCCs during tumor initiation in a manner important for tumor formation in animals (Sendoel et al. 2017).

Recently, eIF2A knock-out mice were found to exhibit reduced longevity, dysregulated lipid metabolism, obesity, decreased glucose tolerance and increased insulin resistance—all phenotypes of metabolic syndrome—and also decreased lymphocyte production and compromised immune tolerance, implicating eIF2A in several important physiological processes and disease states. The mice lacking eIF2A, however, did not exhibit defects in expression of BiP or a protein (CHOP) up-regulated by eIF2α phosphorylation during ER stress (Anderson et al. 2021). It is unknown whether any of the abnormalities observed in eIF2A-deficient mice result from dysregulated translation, nor whether any of the translational functions ascribed to eIF2A in cultured mammalian cells operate *in vivo*.

The amino acid sequence of eIF2A is fairly well conserved between yeast and mammals across the length of the protein (Komar and Merrick 2020), raising the possibility that it might function in IRES-mediated translation or non-canonical initiation at NCC codons in yeast. Analysis of a yeast mutant lacking the gene encoding eIF2A (*YGR054W*) indicated, as might be expected, that eIF2A did not participate in canonical, cap-dependent translation initiation, having no impact on bulk translation initiation *in vivo.* Yeast eIF2A was found however, to co-fractionate with 40S and 80S ribosomes and to interact genetically with canonical initiation factors eIF4E and eIF5B (Zoll et al. 2002; Komar et al. 2005). Surprisingly, analysis of the deletion mutant (referred to below as *eIF2AΔ*) indicated that eIF2A functions to suppress, rather than enhance, non-canonical initiation events that occur independently of scanning, including an IRES identified in the *URE2* mRNA ((Komar et al. 2003; Komar et al. 2005; Reineke and Merrick 2009); reviewed in (Komar and Merrick 2020)). It is unknown whether yeast eIF2A can substitute for eIF2 in Met-tRNAi recruitment under conditions of eIF2α phosphorylation. One piece of evidence arguing against this idea is that the *eIF2AΔ* mutation did not alter expression of a reporter for the *GCN4* transcript (Zoll et al. 2002) that, as mentioned above, is induced by eIF2α phosphorylation. As translational control of *GCN4* mRNA is highly specialized (Gunisova et al. 2018), it seemed possible that other yeast mRNAs might utilize eIF2A as an auxiliary factor for Met-tRNAi recruitment to AUG codons, or for initiation at near-cognate codons. In this study, we explored this possibility by conducting ribosome profiling of an *eIF2AΔ* mutant in the presence or absence of increased phosphorylation of eIF2α induced by amino acid limitation. Our results do not support the possibility that eIF2A frequently substitutes for eIF2 or participates in non-canonical initiation events, even under stress conditions of attenuated eIF2 function.

## RESULTS

### Eliminating yeast eIF2A has little impact on translational reprogramming conferred by phosphorylation of eIF2α in cells starved for amino acids

To examine whether eIF2A provides an eIF2-independent initiation mechanism for any yeast mRNAs, we first examined bulk polysome formation in a yeast mutant lacking the gene *YGR054W* encoding eIF2A (denoted *eIF2AΔ* below) and an isogenic wild-type strain (WT), both grown in nutrient-replete medium (SC) or under conditions of isoleucine/valine starvation, imposed by the drug sulfometuron methyl (SM), to induce eIF2α phosphorylation by Gcn2 and thereby reduce TC levels. We reasoned that if eIF2A can substitute for eIF2 to maintain translation initiation of a sizeable fraction of mRNAs when eIF2 function is reduced, then we might observe a depletion of polysomes in the SM-treated *eIF2AΔ* mutant compared to SM-treated WT cells. As we reported previously (Gaikwad et al. 2021), SM treatment of WT evoked a small reduction in the ratio of polysomes to monosomes (P/M), indicating a diminished rate of bulk translation initiation (Figure 1A(i) & (iii)). Essentially identical P/M ratios were observed in the corresponding untreated and SM-treated *eIF2AΔ* mutant (Figure 1A(ii) & (iv)), suggesting that eIF2A does not function broadly to compensate for reduced eIF2 function in the yeast translatome. These findings are consistent with a previous analysis in which eIF2A was depleted by transcriptional shut-off of an *eIF2A* allele expressed from the *GAL1* promoter (Zoll et al. 2002).

**Figure 1.**
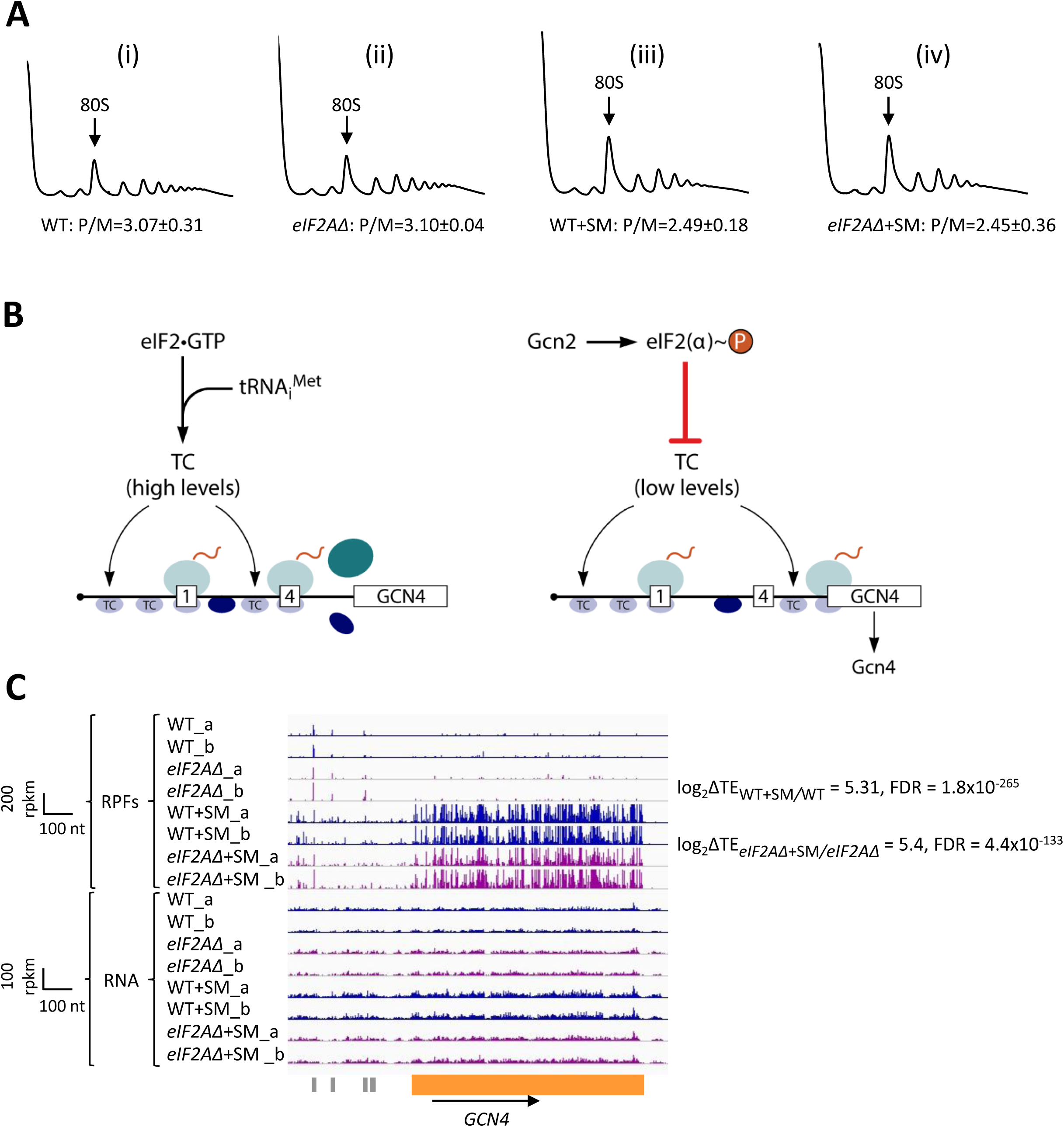
Elimination eIF2A has no effect on bulk protein synthesis in the presence or absence of amino acid starvation. (**A)** Polysome profiles of WT strain (BY4741) and *eIF2AD* mutant (F2247) untreated (i)-(ii) or treated with SM (iii)-(iv). For (i)-(ii), cells were cultured in SC medium at 30°C to log-phase and treated with 50 μg/mL of cycloheximide 5 min prior to harvesting. For (iii)-(iv), cells were cultured in SC medium lacking Ile/Val and treated with 1 µg/mL of SM for 20 min before addition of cycloheximide. Cell extracts were resolved by sedimentation through sucrose density gradients and scanned continuously at 260 nm during fractionation. The plots show the A_260_ measured across the gradient with the top of the gradient on the left. **(B)** Schema of translational control of *GCN4* mRNA, wherein translation of the main CDS is induced by phosphorylation of eIF2α through a specialized “delayed reinitiation” process mediated by four short upstream open reading frames (uORFs). (See text for details). **(C)** Genome browser view of ribosome profiling data for *GCN4* mRNA. Tracks display RPF or mRNA reads mapped across the transcription unit, with the scales given in rpkm (Reads Per Kilobase of transcript per Million mapped reads). Data are presented for WT (blue) and *eIF2Δ* cells (purple) with or without SM treatment, as indicated. Each genotype/treatment includes two biological replicates, designated _a and _b. The main CDS is shown schematically in orange below the tracks; the four uORFs are in grey. The calculated values for log_2_ΔTE_WT+SM/WT_ and log_2_ΔTE*_eIF2A_*_Δ+SM/*eIF2A*Δ+SM_ and the respective FDRs are shown on the right.

We next asked whether eIF2A can provide an eIF2-independent initiation mechanism for any particular mRNAs in yeast, by conducting ribosome profiling of the same *eIF2AΔ* mutant and WT strains under the conditions of SM treatment employed above. Ribosome profiling entails deep-sequencing of 80S ribosome-protected mRNA fragments (RPFs, or ribosome footprints) in parallel with total RNA. The ratio of sequencing reads of RPFs summed over the CDS to the total mRNA reads for the corresponding transcript provides a measure of translational efficiency (TE) for each mRNA (Ingolia et al. 2009). Owing to normalization for total read number in each library, the RPF and mRNA reads and the calculated TEs are determined relative to the average values for each strain. The RPF and RNA read counts between biological replicates for each strain and condition were highly reproducible (Pearson’s r ≈ 0.99) (Figure 1-figure supplement 1A-H).

To establish that SM treatment induced comparable levels of eIF2α phosphorylation in both the WT and *eIF2AΔ* mutant, we examined the ribosome profiling data for *GCN4* mRNA, whose translation is induced by phosphorylated eIF2α by the specialized “delayed reinitiation” mechanism imposed by the four upstream open reading frames in its transcript. Translation of the 5’-proximal uORFs (uORF1 and uORF2) gives rise to 40S subunits that escape recycling at these uORF stop codons and resume scanning downstream. At high TC levels, they quickly rebind TC and reinitiate at uORFs 3 or 4 and are efficiently recycled from the mRNA following their translation. When TC levels are reduced by eIF2α phosphorylation, a fraction of scanning 40S subunits fails to rebind TC until after bypassing uORFs 3-4, and then rebind the TC in time to reinitiate at the *GCN4* CDS, inducing *GCN4* translation (Figure 1B) (reviewed in (Hinnebusch 2005; Gunisova et al. 2018)).

Ribosome profiling revealed the expected strong induction of *GCN4* translation evoked by SM treatment of WT cells, as revealed by greatly increased RPFs in the CDS with little or no change in *GCN4* mRNA reads, yielding an increase in TE (ΔTE) of ∼40-fold (Figure 1C, WT+SM versus WT, cf. replicate cultures _a and _b for RPFs and RNA reads). Highly similar results were observed in the *eIF2AΔ* mutant in the presence and absence of SM (Figure 1C, *eIF2AΔ*+SM versus *eIF2AΔ*), indicating that eIF2α phosphorylation was induced by SM at comparable levels in both the mutant and WT strains. These data argue against the possibility that eIF2A contributes to recruitment of Met-tRNA_i_ as an alternative to the TC by 40S subunits scanning downstream from *GCN4* uORFs 1-2. If this was the case, then the *eIF2AΔ* mutation would impose a further delay in rebinding Met-tRNA_i_ to the 40S subunits re-scanning downstream from uORFs 1-2 and thereby increase the proportion that bypass uORFs 3-4 when TCs are limiting, derepressing *GCN4* translation in SM-treated *eIF2AΔ* versus WT cells. The highly similar induction of *GCN4* translation in the two strains (Figure 1C) argues against such a role for eIF2A in recruiting Met-tRNAi to 40S subunits scanning the *GCN4* mRNA leader. These findings are in agreement with measurements of *GCN4-lacZ* reporter expression in SM-treated *eIF2AΔ* versus WT cells; although the induction by SM was quite modest in that study, and an auxiliary role for eIF2A might have gone undetected (Zoll et al. 2002).

We turned next to the question of whether eliminating eIF2A alters the translation of any other yeast mRNAs, reasoning that mRNAs able to utilize eIF2A in place of TC for recruitment of Met-tRNA_i_ would exhibit greater TE reductions evoked by SM in *eIF2AΔ* mutant versus WT cells. If such mRNAs could be translated efficiently utilizing TC alone, then they would require eIF2A conditionally, i.e., only when eIF2 function is reduced during starvation. If instead such mRNAs rely primarily on eIF2A for efficient initiation regardless of TC levels, they would exhibit reduced TEs in *eIF2AΔ* versus WT cells in the absence of SM treatment. Importantly, DESeq2 analysis of the ribosome profiling data obtained for the *eIF2AΔ* mutant and WT strains cultured in the absence of SM revealed no significant TE reductions in the mutant, as no mRNAs exhibited TE*_eIF2AΔ_*/TE_WT_ ratios < 1 at a false discovery rate (FDR < 0.25) that is appropriate for two highly correlated biological replicates (Lamarre et al. 2018) (Figure 2A). This finding suggests that few, if any, mRNAs are appreciably dependent on eIF2A for translation initiation in nutrient-replete cells where TC is abundant.

**Figure 2.**
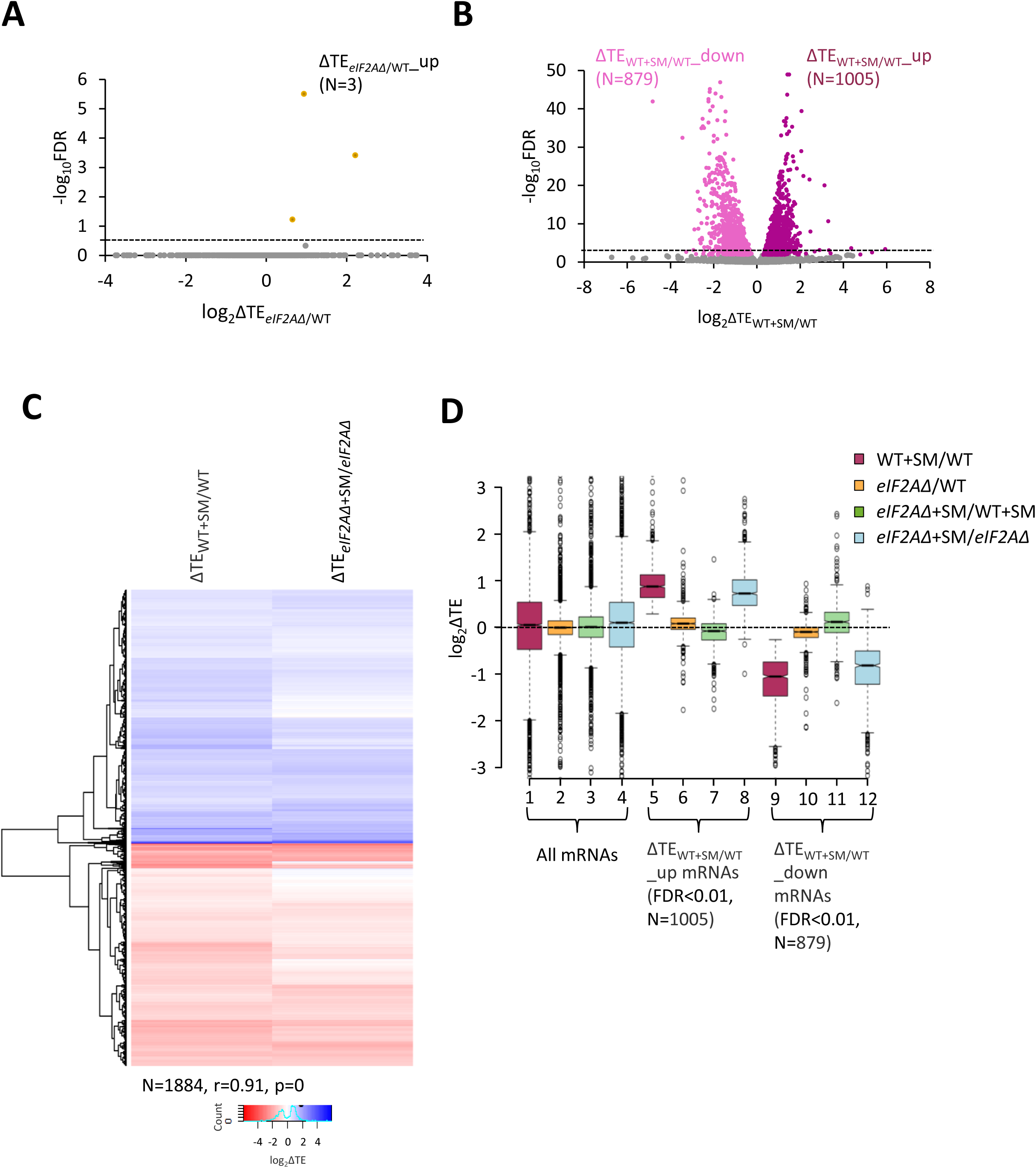
eIF2A is not critical for translation of any individual mRNAs in non-starved cells and has little impact on the reprogramming of TEs conferred by amino acid starvation. **(A)** Volcano plot depicting the log_2_ ratios of TEs in *eIF2AD* versus WT cells (log_2_ΔTE*_eIF2AΔ_*_/WT_ values) for each mRNA (x-axis) versus negative log_10_ of the FDR (y-axis) determined by DESeq2 analysis of ribosome profiling data for the 5340 mRNAs with evidence of translation. Genes showing a significant increase in TE in *eIF2AD* versus WT cells at FDR < 0.25 (ΔTE*_eIF2AΔ_*_+SM/WT__up) are plotted in orange circles. The dotted line marks the 25% FDR threshold, below which all other 5337 mRNAs are plotted in grey. **(B)** Volcano plot as in (A) showing the log_2_ ratios of TEs in WT+SM cells versus WT cells (log_2_ΔTE_WT+SM/WT_ values) for the 5441 mRNAs with evidence of translation. The dotted line marks the 1% FDR threshold. Genes showing a significant increase (ΔTE*_eIF2AΔ_*_+SM/WT__up) or decrease (ΔTE_WT+SM/WT__down) in TE in WT+SM versus WT cells at FDR < 0.01, are plotted in magenta and pink circles, respectively. **(C)** Hierarchical clustering analysis of log_2_ΔTE values for the 1884 mRNAs (arrayed from top to bottom) that exhibit significant TE decreases or increases in SM-treated versus untreated WT cells at FDR < 0.01 (defined in (B)) conferred by SM treatment of WT cells (col. 1) or SM treatment of *eIF2AD* cells (col. 2), with the log_2_ΔTE values represented on a color scale ranging from 4 (dark blue) to -4 (dark red). The Pearson coefficient (r) and corresponding p-value for the correlation between log_2_ΔTE values in the two cols. are indicated below. **(D)** Notched box plots of log_2_ΔTE values for the indicated mutant/condition for all mRNAs (cols. 1-4) or for the indicated mRNA groups identified in (B). The y-axis scale was expanded by excluding a few outliers from the plots.

In addition to induction of *GCN4* translation, SM-treatment of WT cells leads to a broad reprogramming of translational efficiencies, as DESeq2 analysis revealed hundreds of mRNAs exhibiting increases or decreases in relative TE even at the highly stringent FDR of < 0.01 (Figure 2B). Previously, we presented evidence that many of these TE changes conform to a pattern in which mRNAs that are efficiently translated in untreated WT cells tend to exhibit increased relative TEs, whereas poorly translated mRNAs tend to show reduced relative TEs, when phosphorylation of eIF2α is induced by SM. This same pattern was evident under two other conditions in which 43S PIC assembly is reduced, impaired recycling of 40S subunits from post-termination complexes at stop codons and depletion of an essential 40S ribosomal protein. We proposed that this stereotypical reprogramming of translation arises from increased competition among mRNAs for limiting PICs wherein strongly translated mRNAs outcompete weakly translated mRNAs to skew TE increases towards “strong” mRNAs (Gaikwad et al. 2021). SM-treatment of the *eIF2AΔ* mutant also produced a broad reprogramming of TEs involving hundreds of mRNAs translated relatively better or worse on SM treatment (Figure 2-figure supplement 1A). Comparing the TE changes conferred by SM in WT versus *eIF2AΔ* cells for the mRNAs showing TE changes in WT revealed that the majority of transcripts showed TE changes in the same direction on SM treatment of WT and *eIF2AΔ* cells (Figure 2C). Indeed, a very strong positive correlation exists, with a coefficient of 0.91, between the TE changes conferred by SM in the two strains (Figure 2C and Figure 2-figure supplement 1B), thus indicating that elimination of eIF2A did not substantially alter the global reprogramming of translation produced by phosphorylation of eIF2α.

If certain mRNAs depend more heavily on eIF2A for Met-tRNAi recruitment when eIF2α is phosphorylated, we might expect to observe additive reductions in TE when combining elimination of eIF2A by the *eIF2AΔ* mutation with inhibition of eIF2 by SM treatment. Interrogating the large group of 879 mRNAs that showed TE reductions on SM treatment of WT cells revealed that they show a relatively smaller, not larger, decrease in median TE on SM treatment of the *eIF2AΔ* mutant versus SM treatment of WT (Figure 2D, col. 12 versus. col. 9). Consistent with this finding, most of these mRNAs exhibit a small increase in TE on comparing SM-treated *eIF2AΔ* to SM-treated WT cells (Figure 2D, col. 11). (In these and all other box plots, when the notches of different boxes do not overlap, their median values are judged to differ significantly with a 95% confidence level. As shown in cols. 1-4, the median TE change for all ∼5500 expressed mRNAs detected in our profiling experiments is close to unity (log_2_ = 0) for all of the comparisons examined in Figure 2D). These results suggest that the majority of mRNAs whose translation is diminished by eIF2α phosphorylation in WT cells do not utilize eIF2A for a back-up initiation mechanism that would mitigate their TE reductions when eIF2 is impaired. Our finding that most mRNAs exhibit somewhat greater TEs in *eIF2AΔ* versus WT cells when both are treated with SM (Figure 2D, col. 11) might indicate that eIF2A generally acts to repress the translation of these mRNAs rather than augmenting eIF2 function in Met-tRNAi recruitment. This would not be the case in the absence of SM, however, as the results in col. 10 of Figure 2D suggest a small positive effect of eIF2A on translation of these mRNAs in non-starved cells.

### Only few mRNAs exhibit translational reprogramming consistent with eIF2A functioning as a back- up to eIF2

To determine whether there are any individual mRNAs that show greater TE reductions in response to SM treatment when eIF2A is absent, we conducted DESeq2 analysis of the TE changes in SM-treated *eIF2AΔ* versus SM-treated WT cells. A group of only 32 mRNAs showed significant TE reductions in this comparison, i.e., TE*_eIF2AΔ_*_+SM_/TE_WT+SM_ < 1, FDR < 0.25) (Figure 3A). The TE reductions for this group of transcripts (designated ΔTE*_eIF2AΔ_*_+SM/WT+SM__down) in comparison to all mRNAs were nearly 2-fold greater in SM-treated *eIF2AΔ* versus SM-treated WT cells (Figure 3B, col. 1), which are the results expected if they utilize eIF2A as a back-up when eIF2 is impaired by phosphorylation. They also showed TE reductions in SM-treated versus untreated *eIF2AΔ* cells, albeit of lesser magnitude (Figure 3B, col. 2), also in the manner expected if eIF2 and eIF2A play redundant roles in their translation. Functional redundancy is further supported by the findings that both SM treatment of WT cells and elimination of eIF2A from untreated cells does not reduce their median TEs (Figure 3B, cols. 3-4), as only one of the two factors is impaired or eliminated in these latter comparisons. The fact that the median TE of these mRNAs increases rather than decreases on SM treatment of WT (Figure 3B, col. 3) might be explained by proposing that their ability to rely on eIF2A provides them with a competitive advantage with mRNAs that depend solely on eIF2 when the latter is impaired by phosphorylation.

**Figure 3.**
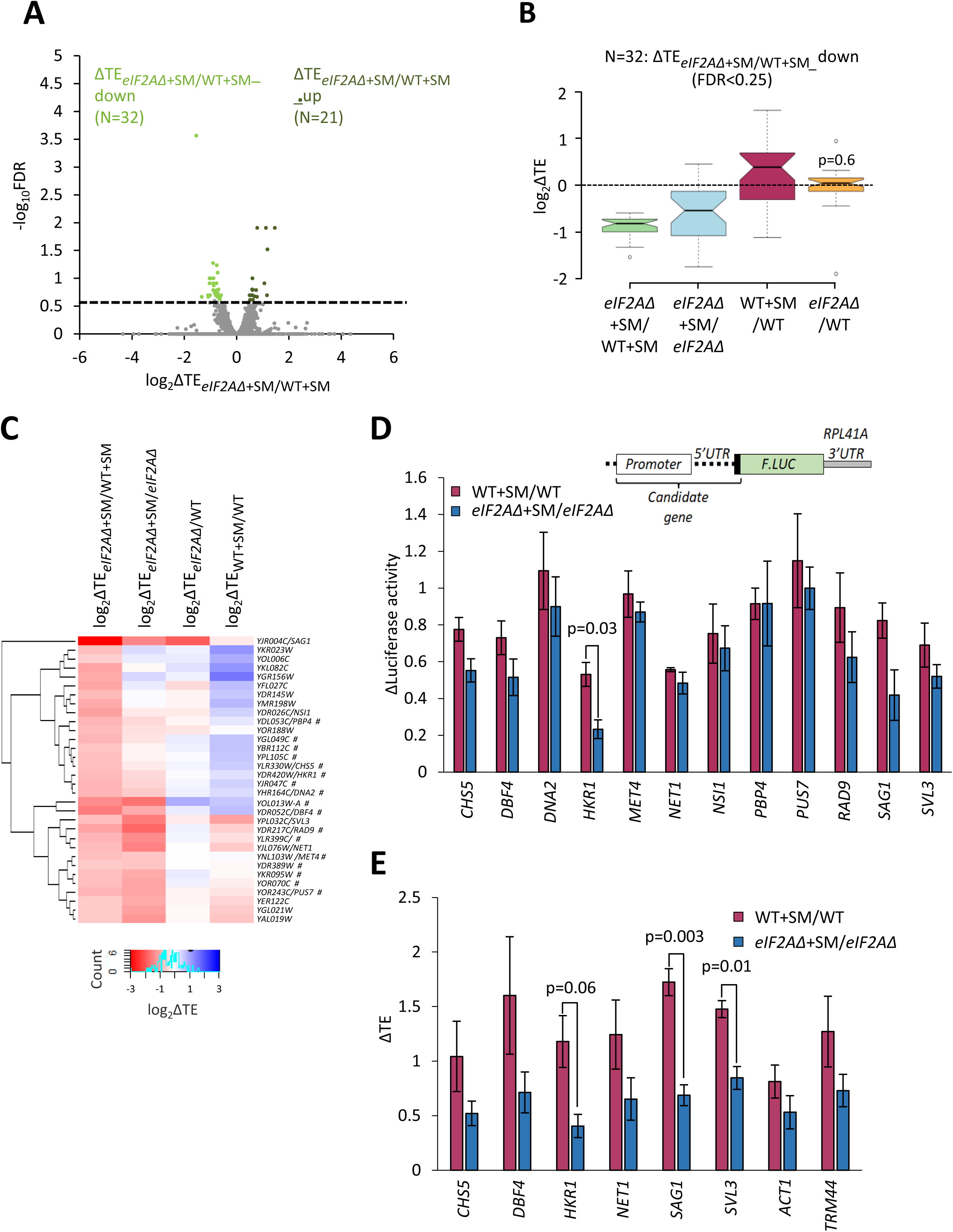
Examination of a small group mRNAs showing evidence of a conditional requirement for eIF2A when eIF2 is impaired. **(A)** Volcano plot as in Figure 2A showing the log_2_ ratios of TEs in *eIF2AD* cells treated with SM versus WT cells treated with SM (log_2_ΔTE*_eIF2AΔ_*_+SM/WT+SM_ values) for the 5482 mRNAs with evidence of translation. The dotted line marks the 25% FDR threshold. Genes exhibiting a significant increase (ΔTE*_eIF2AΔ_*_+SM/WT+SM__up) or decrease (ΔTE*_eIF2AΔ_*_+SM/WT+SM__down) at FDR < 0.25 are plotted in dark or light green circles, respectively. **(B)** Notched box plots of log_2_ΔTE values for the indicated mutant/condition for the 32 mRNAs in the group ΔTE*_eIF2AΔ_*_+SM/WT+SM_down_ defined in (A). The y-axis scale was expanded by excluding a few outliers from the plots. Statistical significance determined using the Mann-Whitney U test is indicated for the changes in col. 4 compared to the changes observed for all mRNAs. **(C)** Hierarchical clustering analysis of log_2_ΔTE values for the 32 mRNAs (arrayed from top to bottom) in the group defined in (A) for the four comparisons listed across the top, with log_2_ΔTE values represented on a color scale ranging from 4 (dark blue) to -4 (dark red). The systematic gene names are listed for all 32 mRNAs, and the common name is indicated for those genes subjected to *LUC* reporter analysis below. Genes marked with “#”s display the pattern of TE changes consistent with conditional stimulation by eIF2A when eIF2 function is reduced by phosphorylation, showing Only 17 of the 32 transcripts (marked with “#”) displayed the diagnostic pattern of an appreciable reduction in TE both on elimination of eIF2A in SM-treated cells and on SM-treatment of cells lacking eIF2A (red or pink hues in cols. 1-2) but either a lesser reduction, no change, or increase in TE on SM-treatment of WT cells and on elimination of eIF2A from untreated cells (light pink, white or blue hues in cols. 3-4). **(D)** Expression of *LUC* reporters in different strains/conditions constructed for selected candidate genes analyzed in (C). The schematic depicts reporter construct design wherein the native gene promoter, 5’ UTR, and first 20 codons of the CDS are fused to firefly luciferase coding sequences (*F.LUC*), followed by a modified *RPL41A* 3’ UTR. Plasmid-borne reporter constructs were introduced into the WT and *eIF2AD* strains and three independent transformants were cultured in SC-Ura medium at 30°C to log phase (-SM) or treated with SM at 1 μg/mL after log-phase growth in SC-Ura/Ile/Val and cultured for an additional 6 h before harvesting. Luciferase activities were quantified in WCEs, normalized to total protein, and reported as fold change in relative light units (RLUs) per mg of protein, as means (±SEM) determined from the replicate transformants. The changes in luciferase activity plotted for each of the two comparisons depicted in the histogram were calculated as ratios of the appropriate mean activities. Results of student’s t-tests of the differences in fold changes between the indicated mutations/conditions are indicated. **(E)** Determination of relative TEs for the native mRNAs of selected candidate genes analyzed in (C & D. Cells were cultured in the four conditions described in (D) and WCEs were resolved by sedimentation through 10%-50% sucrose gradients and fractions were collected while scanning at 260 nm. Total RNA was extracted from 80S and polysome fractions, and the abundance of each target mRNA was quantified in each fraction by qRT-PCR, and normalized for (i) the amounts of 18S rRNA quantified for the same fractions and (ii) for the total amounts of monosomes/polysomes recovered in the gradient. The resulting normalized amounts of mRNA in each fraction were multiplied by the number of ribosomes per mRNA in that fraction, summed across all fractions, and divided by the input amount of mRNA in the WCEs, normalized to *ACT1* mRNA, to yield the TEs for that mRNAs in each condition. (See Methods for further details.) The changes in TE conferred by SM treatment of WT or *eIF2AD* cells were calculated for each replicate culture, untreated of SM-treated, and the mean TE changes with SEMs were plotted for the indicated comparisons. The results of student’s t-tests of the differences in mean TE changes are indicated.

To determine how many of these 32 transcripts exhibit TE reductions exclusively when both eIF2 and eIF2A are impaired/eliminated, we conducted hierarchical clustering of the TE changes in the four comparisons described above, displaying the magnitude of changes with a heat map. Only 17 of the 32 transcripts (marked with “#”) displayed the diagnostic pattern of an appreciable reduction in TE both on elimination of eIF2A in SM-treated cells and on SM-treatment of cells lacking eIF2A (red or pink hues in cols. 1-2 of Figure 3C), but either a lesser reduction, no change, or increase in TE on SM-treatment of WT cells and on elimination of eIF2A from untreated cells (light pink, white or blue hues in cols. 3-4 of Figure 3C).

In an effort to provide independent evidence that a subset of the transcripts analyzed in Figure 3C are dependent on eIF2A only when eIF2 function is diminished by SM, we constructed luciferase reporters for 8 of the aforementioned 17 genes (*CHS5, DBF4, DNA2, HKR1, MET4, PBP4, PUS7, RAD9*) and for 4 other genes that satisfied only one of the two criteria for conditional dependence on eIF2A when eIF2 is impaired by phosphorylation *(SAG1, SVL3, NET1,* and *NSI1).* The promoters and 5’UTR sequences of the candidate genes were fused to the firefly luciferase CDS and the 3’UTR and transcription termination sequences of the yeast *RPL41A* mRNA and introduced into yeast on single-copy plasmids (Sen et al. 2015). Assaying these *LUC* reporters in both WT and *eIF2AΔ* cells in the presence and absence of SM showed that only the *HKR1* reporter displayed a significantly greater reduction in expression in response to SM treatment in *eIF2AΔ* versus WT cells (Figure 3D). Hence, except for *HKR1* mRNA, the *LUC* reporter analysis failed to support a conditional requirement for eIF2A for the candidate genes examined.

As an independent approach, we determined the distributions of native candidate mRNAs that co-sedimented with 80S monosomes or different polysomal species in cell extracts, examining *HKR1* and two other mRNAs (*CHS5* and *DBF4*) that satisfied both of the aforementioned criteria for conditional dependence on eIF2A, three mRNAs (*NET1, SAG1, SVL3*) that satisfied only one of the two criteria, and *ACT1* and *TRM44* mRNAs examined as negative controls. The amounts of each mRNA found in monosome or polysomal fractions was multiplied by the number of ribosomes per mRNA in that fraction (1 for monosomes, 2 for disomes, 3 for trisomes, etc.) to calculate the abundance of ribosomes translating the mRNA and normalized to the input level of the mRNA in the unfractionated extracts to calculate the translational efficiency (TE) of the transcript in each condition. Analyzing the results from three biological replicates for WT versus *eIF2AΔ* cells +/- SM revealed that only three transcripts, *HKR1, SAG1,* and *SVL3,* displayed a greater TE reduction in response to SM treatment of *eIF2AΔ* cells compared to SM treatment of WT (Figure 3E & Figure 3-figure supplement 1). Thus, only the *HKR1* transcript satisfied all of the criteria for conditional eIF2A dependence in ribosome profiling data in a manner confirmed by both reporter analysis and polysome profiling. We conclude that there are very few mRNAs, possibly only one, that utilize eIF2A as a back-up for eIF2 in recruitment of tRNAi when eIF2 is impaired by phosphorylation.

### Yeast eIF2A does not appear to modulate IRES activity even when eIF2 is impaired

We next interrogated our ribosome profiling data for the effects of deleting *eIF2A* on translation of *URE2* mRNA, reported to contain an internal ribosome entry site (IRES) that is inhibited by eIF2A (Komar et al. 2005). Surprisingly, we found no significant change in TE or ribosome occupancy for *URE2* mRNA on deletion of *eIF2A* in either non-starved or SM-starved cells (Figure 4A), providing no evidence that eIF2A acts to repress *URE2* translation under the conditions of our experiments. It was reported that yeast mRNAs *GIC1* and *PAB1* also contain IRESs and are subject to translational repression by eIF2A (Reineke and Merrick 2009), however, we found no significant alteration in their TEs between *eIF2AΔ* and WT cells in the presence or absence of SM treatment (Figure 4B-C). Our results do not support a role for eIF2A in repressing IRES function in yeast cells.

**Figure 4.**
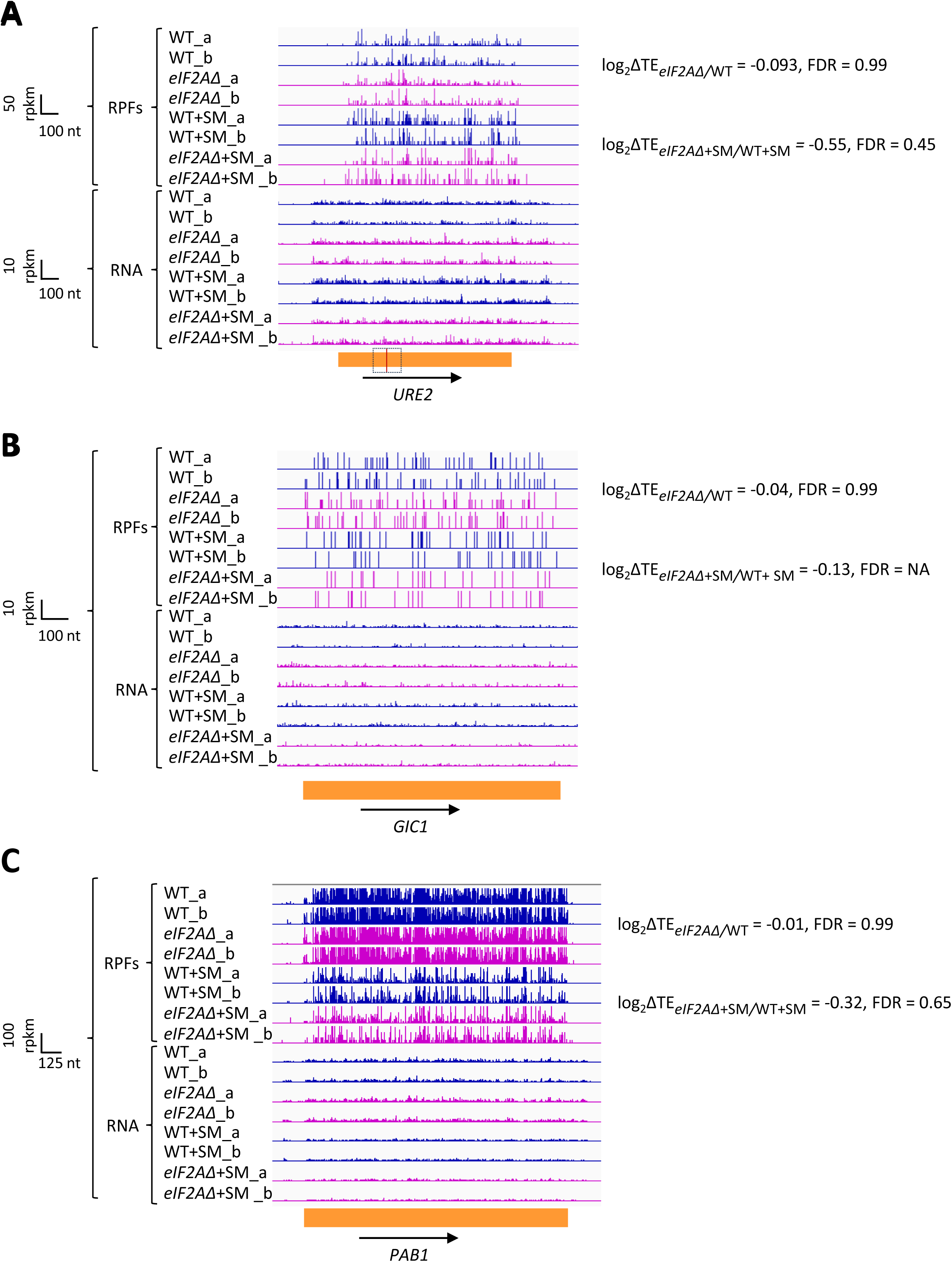
eIF2A has little or no effect on the translation of three mRNAs reported to contain IRESs. Genome browser views of RPF and RNA reads from ribosome profiling data for **(A)** *URE2* mRNA **(B)** *GIC1* mRNA and **(C)** *PAB1* mRNA presented as in Figure 1C. The calculated values for log_2_ΔTE*_eIF2A_*_Δ/WT_ and log_2_ΔTE*_eIF2A_*_Δ+SM/WT+SM_ with the respective FDRs are shown on the right. The scale is shown on the left. The region containing the *URE2* IRES is enclosed in a dotted box, with the AUG start codon highlighted in red. Locations of the *GIC1* and *PAB1* IRESs have not been defined.

### Eliminating eIF2A appears to have little consequence on translation of upstream open-reading frames

It has been reported that mammalian eIF2A is required for the translation of certain non-AUG initiated uORFs and that translation of these uORFs upregulates translation of the downstream CDS when eIF2α is phosphorylated during ER stress (Starck et al. 2016) or during tumor initiation (Sendoel et al. 2017). Hence, we examined our profiling data more closely to determine if eIF2A might affect the translation of mRNAs harboring uORFs in yeast cells. We did not restrict our attention to stimulatory uORFs, which are rare in yeast (May et al. 2023), but included all uORFs in our analyses.

In addition to *GCN4* described above, *CPA1* mRNA, encoding an arginine biosynthetic enzyme, contains a single AUG-initiated inhibitory uORF that attenuates translation of the main CDS in cells replete with arginine (Werner et al. 1987; Delbecq et al. 1994). In accordance with previous findings that the inhibitory effects of single uORFs can be diminished by eIF2α phosphorylation by allowing scanning ribosomes to bypass the uORF start codon (Young and Wek 2016; Dever et al. 2023), we observed a ∼2-fold increase in the TE of *CPA1* mRNA on SM-treatment of WT cells (Figure 5A). However, this TE change was not significantly altered by the *eIF2AΔ* mutation. Similarly, for the well-characterized *YAP2/CAD1* mRNA, containing two inhibitory AUG uORFs (Vilela et al. 1998), we observed a 2.6-fold TE increase conferred by SM in WT cells that, again, was not significantly altered by deletion of *eIF2A* (Figure 5B). The *eIF2AΔ* deletion also had no effect on the TEs of *CPA1* and *YAP2/CAD1* in non-starved cells. Thus, although we found evidence that translational repression exerted by the uORFs in *CPA1* and *YAP2/CAD1* mRNAs is mitigated by eIF2α phosphorylation, it appears that eIF2A has little or no role in recognizing the AUG codons of these uORFs in the presence or absence of diminished TC levels, just as we concluded above for the *GCN4* uORFs.

**Figure 5.**
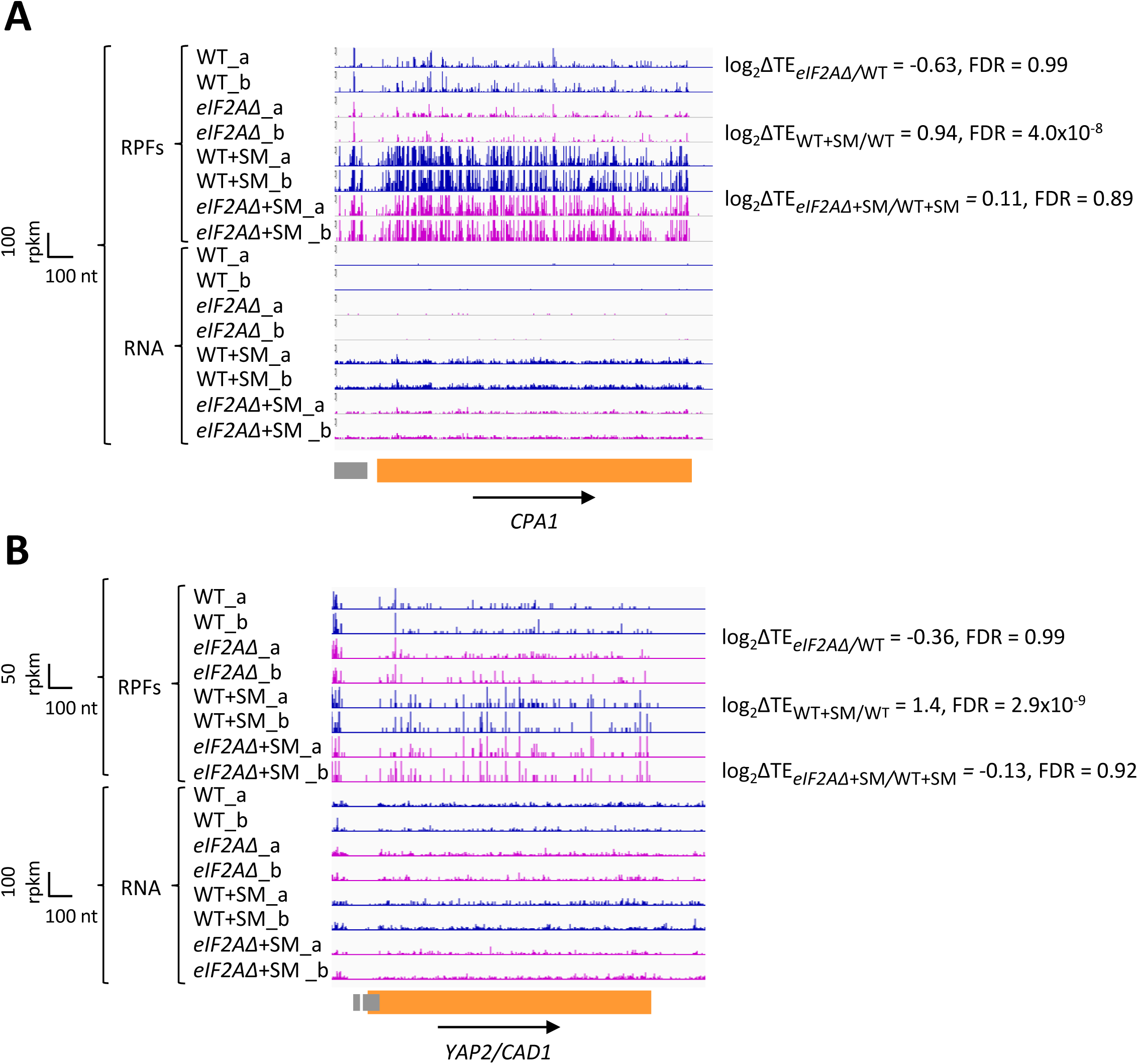
eIF2A plays little or no role in uORF-mediated translational control of *CPA1* or *YAP2/CAD1* mRNA. Genome browser views of RPF and RNA reads from ribosome profiling data for **(A)** *CPA1* mRNA and **(B)** *YAP2/CAD1* mRNA presented as in Figures 1C & 4. CDS and uORFs are represented in orange and grey rectangles, respectively.

Looking more broadly, we interrogated a previously identified group of 1306 mRNAs containing 2720 uORFs, initiating with either AUG or one of the nine NCCs, which showed evidence of translation in multiple ribosome profiling datasets from various mutant and WT strains (Zhou et al. 2020). We also analyzed a second set of 791 mRNAs containing 982 AUG- or NCC-initiated uORFs that are both evolutionarily conserved and show evidence of translation in ribosome profiling experiments (Spealman et al. 2018). These four groups of mRNAs with AUG- or NCC-uORFs showed little or no TE change for their CDSs on SM treatment of WT cells (Figure 6A(i)-(ii), cols. 4 and 7 versus 1), indicating that, in contrast to *GCN4, CPA1,* and *YAP2/CAD1* mRNAs, most of them do not contain inhibitory uORFs that can be bypassed in response to eIF2α phosphorylation. Both groups of mRNAs containing either AUG- or NCC-initiated uORFs exhibit very small, albeit highly significant, reductions in CDS TEs in the *eIF2AΔ* mutant versus WT cells in the absence of SM (Figure 6A(i)-(ii), cols. 5 and 8 versus 2); however, the same was not observed for the effects of *eIF2AΔ* in SM-treated cells where eIF2 function is attenuated (Figure 6A(i)-(ii), cols. 6 and 9 versus 3).

**Figure 6.**
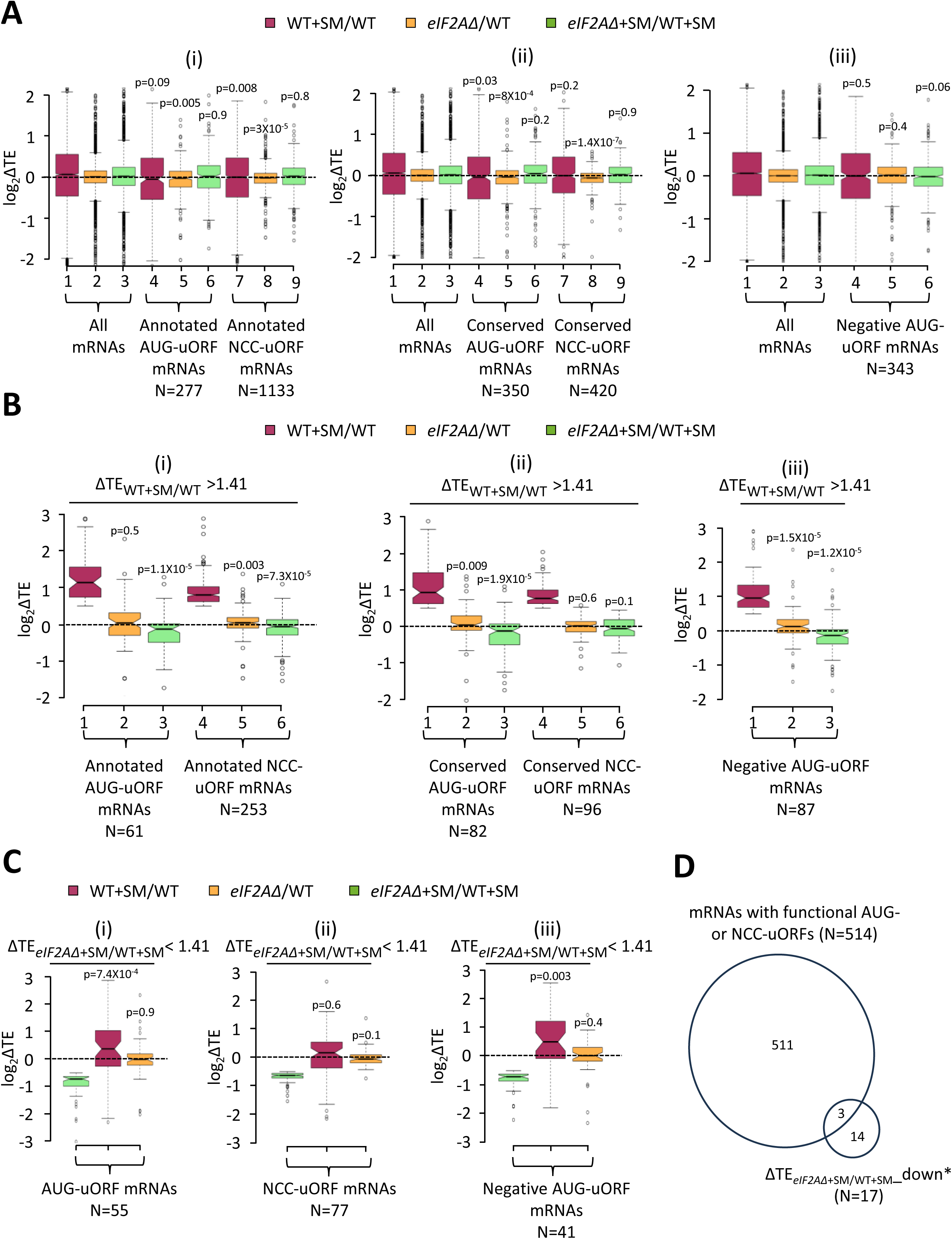
Minimal effects of eliminating eIF2A on translation of mRNAs harboring translated uORFs. **(A)** Notched box plots of log_2_ΔTE values for all mRNAs (for which TEs could be determined from our ribosome profiling data) containing annotated AUG- or NCC-uORFs (i), conserved AUG- or NCC-uORFs (ii), or single functional inhibitory AUG-uORFs (iii), conferred by SM treatment of WT cells (maroon), by the *eIF2AD* mutation in untreated cells (orange), or by the *eIF2AD* mutation in SM-treated cells (green). Statistical significance determined using the Mann-Whitney U test is indicated for selective comparisons of changes observed for the indicated groups in comparison to the changes for all mRNAs. A few outliers were omitted from the plots to expand the y-axis scale. **(B)** Notched box plots as in (A) for the subsets of the same mRNA groups analyzed there exhibiting > 1.41-fold increases in TE in SM-treated versus untreated WT cells. A few outliers were omitted from the plots to expand the y-axis scale. Statistical significance determined as in (A). **(C)** Notched box plots as in (A-B) for the subsets of the mRNA groups analyzed there exhibiting > 1.41-fold decreases in TE in SM-treated *eIF2AΔ* versus SM-treated WT cells. A few outliers were omitted from the plots to expand the y-axis scale. Statistical significance determined as in (A). **(D)** Proportional Venn diagram showing overlap between the 17 mRNAs identified in Figure 3A showing evidence for a conditional requirement for eIF2A when eIF2 function is reduced by SM (transcripts marked with “#”s) and the 514 mRNAs bearing functional AUG or NCC-uORFs.

Evidence of translation of a uORF does not guarantee that it has an appreciable impact on the proportion of scanning ribosomes that reach the downstream CDS. Recently, groups of 557 mRNAs containing AUG-initiated uORFs and 191 mRNAs with NCC-uORFs were identified in which the uORFs were shown to influence translation of downstream CDSs by massively parallel analysis of reporters containing the native 5’UTRs in comparison to mutant reporters lacking the uORF start codons (May et al. 2023). It was reported that among the 407 mRNAs containing a single functional AUG-uORF wherein mutating the uORF significantly altered reporter expression by > 1.5-fold, all but 13 of the uORFs functioned to inhibit reporter expression. Among the 144 mRNAs containing a single functional NCC-uORF, only 13 of the uORFs influenced reporter expression either positively or negatively by > 1.5-fold, reflecting the weaker effects of NCC-versus AUG-initiated uORFs on downstream translation (May et al. 2023). Our analysis of the 394 mRNAs containing a single inhibitory AUG-uORF—the only group large enough for statistical analysis of TE changes—revealed little or no change in median TE in response to SM treatment or to the *eIF2AΔ* mutation in the presence or absence of SM (Figure 6A(iii), cols. 4-6 versus 1-

3). These findings suggest that eIF2A has a minimal role in determining whether functional inhibitory AUG-initiated uORFs are translated or bypassed by scanning ribosomes, in the presence or absence of eIF2α phosphorylation. These results support our findings above that the majority of mRNAs containing translated uORFs show little response to either eIF2α phosphorylation, the elimination of eIF2A, or a combination of both perturbations.

We probed more deeply into the three sets of uORF-containing mRNAs mentioned above by examining the subsets of each group that exhibit > 1.41-fold increases in TE in response to SM treatment of WT cells, making them novel candidates for mRNAs controlled by inhibitory uORFs that can be overcome by eIF2α phosphorylation. These uORF-containing mRNAs exhibit median TE increases of ∼2- fold on SM treatment of WT cells (Figure 6B(i)-(iii), maroon data), similar to the findings above for *CPA1* and *CAD1* mRNAs (Figure 5A-B). Unlike the larger groups of uORF-containing mRNAs from which they derive (analyzed in Figure 6A(i)-(iii)), these subsets showing appreciable TE induction by SM exhibit slightly reduced median TEs in response to the *eIF2AΔ* mutation in SM-treated, but not untreated cells (Figure 6B(i)-(iii), green versus orange data). One way to explain these findings would be to propose that these mRNAs generally contain a positive-acting uORF that functions to overcome the translational barrier imposed by a second inhibitory uORF located further downstream in response to eIF2α phosphorylation, in the manner established for the positive-acting uORF1 in *GCN4* mRNA. The TE reduction conferred by *eIF2AΔ* exclusively in SM-treated cells would then arise from eIF2A-dependent translation of the stimulatory upstream uORFs in these transcripts to enable scanning ribosomes to bypass the downstream inhibitory uORFs when eIF2 is phosphorylated.

To explore this possibility further, we focused on the uORF-containing mRNAs that are the most dependent on eIF2A for their translation in SM-treated cells, showing a > 1.41-fold reduction in TE in SM-treated *eIF2AΔ* versus SM-treated WT cells. Examining all such mRNAs containing either (i) annotated or conserved AUG-uORFs (Figure 6C(i)), (ii) annotated or conserved NCC-uORFs (Fig, 6C(ii)), or (iii) functional inhibitory AUG-uORFs (Figure 6C(iii)), revealed that all three groups show TE reductions in response to the *eIF2AΔ* mutation only in the presence of SM (green versus orange data) and, except for the NCC-uORF mRNAs, also show TE increases in response to SM in WT cells (6C(i)-(iii), maroon data). This is the same pattern we observed for all of the uORF-containing mRNAs showing appreciable TE increases on SM-treatment of WT cells just described (Figure 6B(i)-(iii)). Moreover, this pattern of TE changes was also found for the group of 32 mRNAs showing highly significant TE reductions conferred by *eIF2AΔ* in SM-treated cells versus SM treatment of WT (Figure 3B); although, only 17 of those transcripts conform to this pattern of TE changes (blue in col. 4 but nearly white in col. 3 of Figure 3B; dubbed ΔTE*_eIF2AΔ_*_+SM/WT+SM__down*). Of these 17 mRNAs, only three (*YFL027C, YKL082C,* and *YLR330W/CHS5*) contain a functional AUG- or NCC-initiated uORF (Figure 6D), which does not represent a statistically significant enrichment for such mRNAs (p = 0.21). Moreover, our *LUC* reporter and polysome profiling analyses of one of the three transcripts, *YLR330W/CHS5,* failed to confirm a conditional dependence for eIF2A (Figure 3D-E). The one mRNA for which conditional eIF2A dependence was confirmed, *YDR420W/HKR1,* does not contain a functional or conserved uORF nor even one merely showing evidence of translation in ribosome profiling experiments.

As an orthogonal approach to detecting a role for eIF2A in regulating the translation of negative-acting uORFs, we reasoned that decreased translation of inhibitory uORFs in *eIF2AΔ* cells would increase the translation of downstream CDSs, leading to a decrease in the ratio of RPFs in uORFs relative to RPFs in the CDSs, which we termed relative ribosome occupancy (RRO). To examine this possibility, we employed DESeq2 to identify statistically significant changes in RRO values for the same groups of mRNAs analyzed above in Figure 6A-B containing annotated or conserved uORFs with either AUG- or NCC-start sites, and which also showed evidence of translation in both WT and *eIF2AΔ* cells. We found that no mRNAs exhibited significant changes in RRO in the *eIF2AΔ* mutant versus WT in the absence or presence SM, even at a relatively non-stringent FDR of 0.5 (Figure 6-figure supplement 1A-D). Furthermore, the groups of mRNAs with annotated or conserved AUG- or NCC-initiated uORFs showed little or no difference in median RRO in the *eIF2AΔ* mutant versus WT cells in the absence or presence of SM (Figure 6-figure supplement 1E-H, yellow versus blue data). At least for the annotated uORFs, the median RRO values are generally higher for mRNAs containing AUG-initiated versus NCC-initiated uORFs in the presence or absence of SM (Figure 6-figure supplement 1 E & G, blue data), as would be expected from more efficient initiation at AUG versus NCC start codons. Thus, our results provide no compelling evidence that eIF2A frequently enhances initiation at negative-acting uORFs initiated by either AUG or NCC start codons in either starved or non-starved cells.

### eIF2A does not affect decoding rates at particular codon combinations during translation elongation

Translation elongation factor eIF5A acts broadly to stimulate the rates of decoding and termination, but is particularly important for certain combinations of codons, which include stretches of proline codons and various 3-codon combinations of Pro, Asp, and Gly codons (Saini et al. 2009; Gutierrez et al. 2013; Schuller et al. 2017). Slower decoding rates for these codon combinations were detected by ribosome profiling of yeast cells depleted of eIF5A by computing pause scores for all ∼8,000 three-amino acid motifs (Schuller et al. 2017). The pause score for each motif is calculated from its 80S occupancy relative to the surrounding stretch of coding sequences and averaged across the translatome. Analyzing our profiling data revealed no significant increase in pause scores for any tripeptide motif in the *eIF2AΔ* mutant versus WT cells. As a positive control for the analysis, we examined our data on WT cells treated or untreated with SM, observing a marked increase in pause scores for all tripeptide motifs with valine in the third position, placing the corresponding Val codons in the A-site of the decoding ribosomes (Figure 7B). Increased pausing at these motifs is expected from reduced levels of charged valyl-tRNA on inhibition of valine biosynthesis by SM and attendant slower decoding of all Val codons. It is unclear why increased pausing is not found at codons for isoleucine, as SM inhibits an enzyme common to the Ile and Val biosynthetic pathways (Jia et al. 2000); however, we note that flux through these pathways is differentially regulated and that the Val and leucine pathways compete for a common precursor (Jones and Fink 1982).) Taken together, our findings indicate that eIF2A plays no major role in stimulating translation elongation through particular tripeptide motifs.

**Figure 7.**
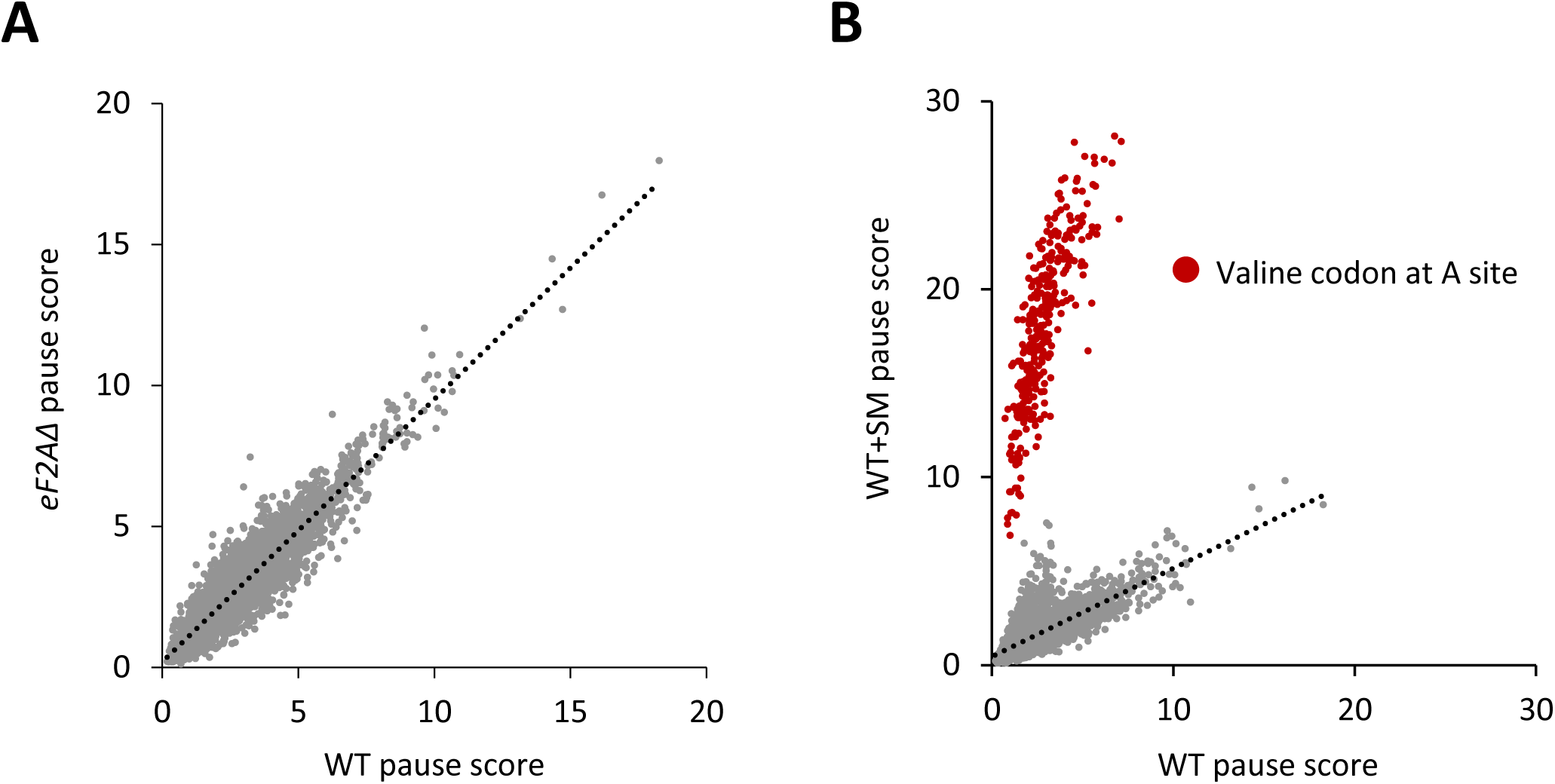
eIF2A plays no major role in stimulating translation elongation for particular tripeptide motifs. **(A)** Scatterplot of average pause scores for 8006 tripeptide motifs, comparing the two biological replicates of ribosome profiling data for *eIF2AΔ* versus WT cells. Each dot on the plot represents a tripeptide motif. Pause scores were computed using a shift value of 18 nt from the 3′-end of the footprint, positioning the first codon of the tripeptide motif in the E site. **(B)** Scatterplot of average pause scores for 6267 tripeptide motifs, comparing the two biological replicates of SM-treated versus untreated WT cells. All 351 detected motifs with valine codons in the A site are highlighted in red. Pause scores were computed as in (A).

### Exploring a possible role for eIF2A in purine biosynthesis

As noted above, a synthetic growth phenotype was observed on combining the *eIF2AΔ* mutation with either a temperature-sensitive mutation affecting eIF4E (Komar et al. 2005) or deletion of the *FUN12* gene encoding eIF5B (Zoll et al. 2002), suggesting functional interactions between eIF2A and these initiation factors. Other genetic interactions involving the *eIF2AΔ* mutation were uncovered in a global analysis of synthetic phenotypes produced by phenotyping double mutants that combine deletions or point mutations in ∼90% of all yeast genes (Costanzo et al. 2016; van Leeuwen et al. 2016), compiled at (https://thecellmap.org/costanzo2016/. The profile of genetic interactions observed for the *eIF2AΔ* mutation was found to significantly resemble that of an *ade1Δ* mutation, including synthetic growth defects in *eIF2AΔ* double mutants containing deletions of genes *HPT1* and *FCY2*, whose products function in purine base and cytosine uptake (Fcy2) or in synthesizing purine nucleotides from purine bases via the salvage pathway (Hpt1) (Ljungdahl and Daignan-Fornier 2012). These findings suggested that eIF2A might function as a positive regulator of de novo purine biosynthesis, such that the *eIF2AΔ* mutation would mimic *adeΔ* mutations in reducing growth in combination with the *hpt1Δ* and *fcy2Δ* mutations as a result of simultaneously impairing both the de novo and salvage pathways for adenine biosynthesis (Ljungdahl and Daignan-Fornier 2012). To test this possibility, we generated the *fcy2ΔeIF2AΔ* double mutant and tested it for growth on synthetic minimal (SD) or synthetic complete (SC) medium supplemented with varying concentrations of adenine. Contrary to expectations, we failed to detect any reduction in growth in the double mutant versus either single mutant or the WT strain in the presence or absence of adenine supplements (Figure 6-figure supplement 2A-B). We also interrogated our ribosome profiling data to examine the effects of the *eIF2AΔ* mutation on expression of the 17 yeast genes encoding proteins involved in de novo synthesis of inosine, adenine or guanine nucleotides (Ljungdahl and Daignan-Fornier 2012). The *eIF2AΔ* mutation conferred no significant reductions in the RPFs, mRNA levels, or TEs of any of 17 genes in non-starvation conditions. Moreover, assaying reporters for two such genes, *IMD2* and *IMD3,* in cells cultured under the conditions of our ribosome profiling experiments revealed no significant reductions in expression conferred by elimination of eIF2A (Figure 6-figure supplement 2C). Our findings do not support the possibility that eIF2A promotes expression of the biosynthetic genes for *de novo* purine biosynthesis.

## DISCUSSION

The mammalian eIF2A protein has been implicated in substituting for eIF2 in recruiting Met-tRNAi to certain viral mRNAs that do not require ribosomal scanning for AUG selection under conditions where eIF2α is being phosphorylated and TC assembly is impaired in virus-infected cells. Other studies have identified a role for eIF2A in initiation at near-cognate initiation codons (Komar and Merrick 2020). As the yeast and mammalian eIF2A proteins show considerable sequence similarity, we looked for evidence that the yeast protein could provide a back-up mechanism for Met-tRNAi recruitment when eIF2α has been phosphorylated by Gcn2 in cells starved for isoleucine and valine using the antimetabolite SM. At an SM concentration adequate to reduce bulk polysome assembly and strongly derepress translation of *GCN4* mRNA in WT cells, neither response was detectably altered by the *eIF2AΔ* mutation. Ribosome profiling revealed a broad reprogramming of translational efficiencies conferred by SM treatment of WT cells, noted previously (Gaikwad et al. 2021), wherein efficiently translated mRNAs tend to show increased relative TEs at the expense of poorly translated mRNAs, which also appeared to be essentially intact in cells lacking eIF2A. Interrogating the TE changes of individual mRNAs, we found only three in the entire translatome that were significantly altered in *eIF2AΔ* cells versus WT in the absence of SM, with all three showing higher, not lower, TEs in the mutant. Thus, we failed to identify even a single mRNA among the 5340 transcripts with evidence of translation that is dependent on eIF2A for efficient translation in non-stressed cells containing normal levels of TC assembly. Under conditions of SM treatment, we identified 32 mRNAs showing significantly reduced TEs in *eIF2AΔ* versus WT cells, consistent with the possibility that they require eIF2A for Met-tRNAi recruitment only when TC assembly is diminished by eIF2α phosphorylation. However, only 17 of these transcripts showed a pattern of TE changes fully consistent with a conditional requirement for eIF2A under conditions of reduced eIF2 function, exhibiting greater TE decreases when both eIF2 function was impaired by phosphorylation and eIF2A was eliminated from cells. Moreover, we could validate this conditional eIF2A dependence by *LUC* reporter analysis and by measuring TE changes of native mRNAs by polysome profiling for only a single mRNA, *HKR1*. A possible limitation of our *LUC* reporter analysis in Figure 3D was the lack of 3’UTR sequences of the cognate genes, which might be required to observe eIF2A dependence. Given that native mRNAs were examined in the orthogonal assay of polysome profiling in Figure 3E, the positive results obtained there for *SAG1* and *SVL3* in addition to *HKR1* should be given greater weight. Nevertheless, our findings indicate a very limited role of yeast eIF2A in providing a back-up mechanism for Met-tRNAi recruitment when eIF2 function is diminished by phosphorylation of its α-subunit.

We also looked intensively at groups of mRNAs containing uORFs to determine if eIF2A might promote uORF translation and thereby influence translation of the downstream CDSs. As observed for *GCN4* mRNA, eliminating eIF2A had no effect on the TEs of *CPA1* and *CAD1* mRNAs, which contain single inhibitory uORFs that appear to be suppressed by eIF2α phosphorylation. We interrogated three different compilations of yeast mRNAs containing translated uORFs containing AUG or NCC start codons, including those with functional evidence of controlling translation of the downstream CDSs or showing evolutionary sequence conservation and, once again, found no evidence for a significant role of eIF2A in either increasing or decreasing the TEs of such transcripts. It remains to be seen if there are any uORFs in yeast mRNAs that depend on eIF2A for their translation. It should also be noted that we could not confirm published evidence indicating that eIF2A inhibits utilization of IRES elements identified previously in *URE2, PAB1,* and *GIC1* mRNAs. Finally, we found no evidence that eIF2A affects the rate of decoding of any particular tri-peptide motifs in the elongation phase of protein synthesis.

The lack of evidence in our study for eIF2A function in translation initiation stands in contrast to findings of negative genetic interactions between the *eIF2AΔ* and mutations in yeast eIF4E (Komar et al. 2005) and eIF5B (Zoll et al. 2002) wherein combining the mutations conferred a more severe cell growth defect. Synthetic negative interactions between *eIF2AΔ* and deletions of each of the two genes encoding eIF4A (*TIF1* and *TIF2*) were also reported in a global analysis of synthetic phenotypes observed in double mutants that combine deletions or point mutations in ∼90% of all yeast genes (Costanzo et al. 2016; van Leeuwen et al. 2016), compiled at (https://thecellmap.org/costanzo2016/). One possibility is that eIF2A is functionally redundant with these initiation factors rather than with eIF2. Another is that eliminating eIF2A affects a different cellular process and that expression of one or more genes involved in that process is diminished by mutations in eIF4E, -4A, and -5B.

A recent study indicated that human recombinant eIF2A inhibits translation in rabbit reticulocyte lysate (RRL) of all mRNAs tested, including one driven by the cricket paralysis virus IGR IRES that requires no canonical initiation factors (Grove et al. 2023). Because the protein binds to 40S subunits and the addition of excess 40S subunits to RRL mitigates its inhibitory effect, it was concluded that human eIF2A sequestered 40S subunits in an inactive complex. While this finding is consistent with the conclusion that yeast eIF2A represses IRES-driven translation in yeast, the finding that *eIF2AΔ* does not increase bulk translation or impact yeast cell growth obtained here and elsewhere (Zoll et al. 2002) suggests that eIF2A does not appreciably sequester 40S ribosomes in yeast cells, even in stress conditions of eIF2α phosphorylation (Figure 1A). The similar finding that knocking out the eIF2A gene had no detectable effect on bulk translation in keratinocytes (Sendoel et al. 2017) led Grove et al. (2023) to surmise that mammalian eIF2A is not normally present in the cytoplasm and would have to be released from a different cellular compartment to impact translation (Grove et al. 2023).

The profile of genetic interactions observed for the *eIF2AΔ* mutation in the aforementioned global analysis of synthetic phenotypes was found to resemble that of an *ade1Δ* mutant (Costanzo et al. 2016; van Leeuwen et al. 2016), lacking an enzyme of *de novo* biosynthesis of AMP and GMP, which suggested that eIF2A might be a positive effector of purine biosynthesis. However, we could not confirm the reported negative genetic interaction between the *eIF2AΔ* and *fcy2Δ* mutations, and our ribosome profiling data revealed no effect of *eIF2AΔ* on the RPF or mRNA abundance of the 17 genes involved in *de novo* purine biosynthesis.

It is surprising that we found so little evidence that eIF2A functions in the yeast translatome even when the activity of the canonical factor eIF2 is attenuated by amino acid starvation. It is possible that eIF2A plays a more important role in translation in some other stress condition, e.g. one in which it is released into the cytoplasm (Grove et al. 2023), or that it functions in a different cellular process altogether in budding yeast. eIF2A may have acquired a new role in controlling translation initiation during mammalian evolution, or it could have lost its function in translation and acquired a different one during fungal evolution.

## MATERIALS AND METHODS

### Yeast Strain and Plasmid Construction

Yeast strains used in this study are listed in Table 1. The primers used for strain construction and verification are listed in Table 3. Strain SGY3 (*eIF2ADfcy2D*) was generated through a two-step process. First, the *kanMX* cassette of the *eIF2AΔ* deletion allele, *ygr054wD::kanMX4,* in strain F2247 was swapped with a hygromycin-resistance cassette to produce SGY1 (*ygr054wD::hphMX4*) by transforming F2247 with a DNA fragment containing the *hphMX4* allele that was PCR-amplified from plasmid p4430. Transformants were selected on YPD agar plates supplemented with 300 μg/mL hygromycin B. In the second step, the *FCY2* gene was deleted in SGY1 by transformation with a DNA fragment containing the *fcy2D::kanMX4* allele PCR-amplified from the genomic DNA of strain F2379. Transformants were selected on YPD agar plates containing 200 μg/mL geneticin (G418) to produce SGY3. The *ygr054wD::hphMX4* and *fcy2D::kanMX4* alleles in strains SGY1 and SGY3, respectively, were verified by PCR analysis of chromosomal DNA using the appropriate primers listed in Table 3.

**Table 1.**
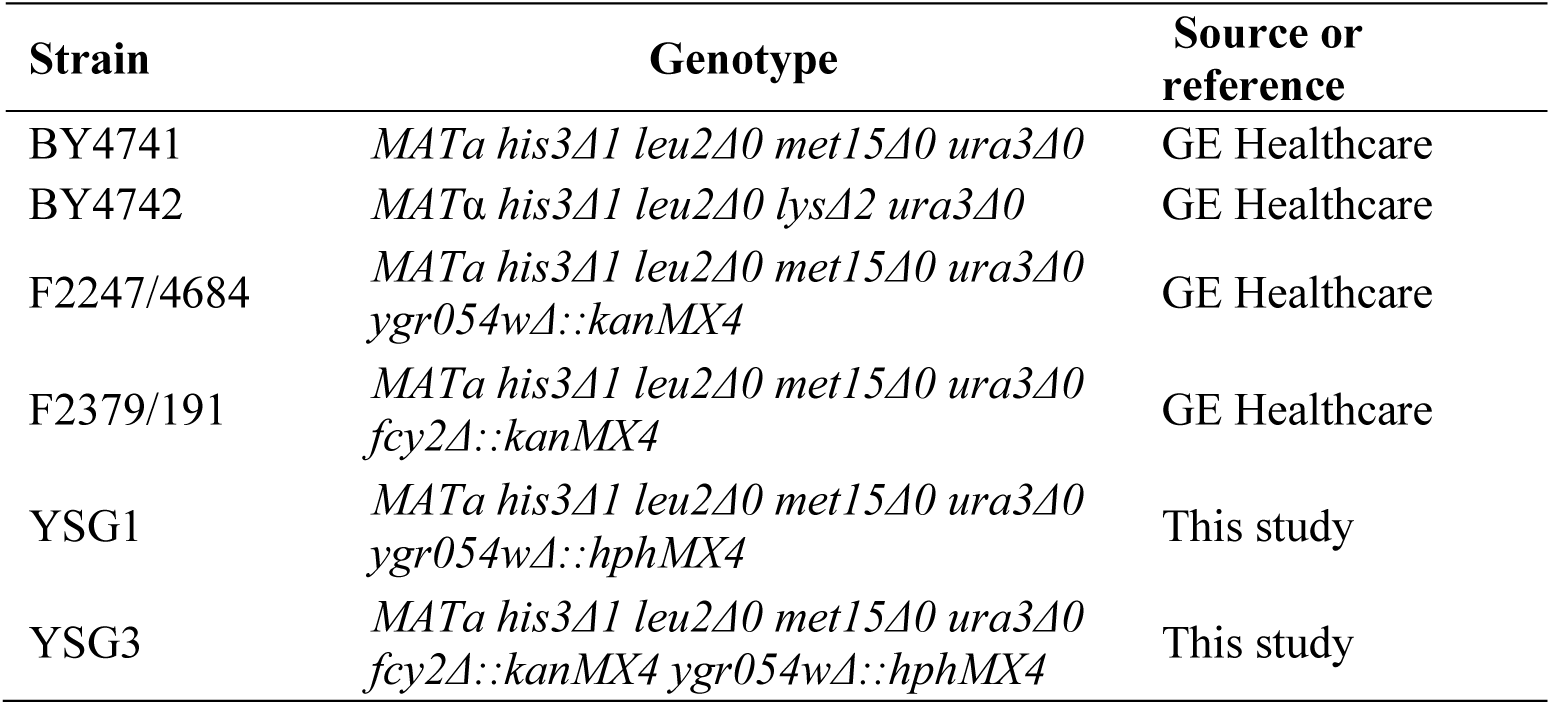
Yeast strains used in this study.

**Table 2.**
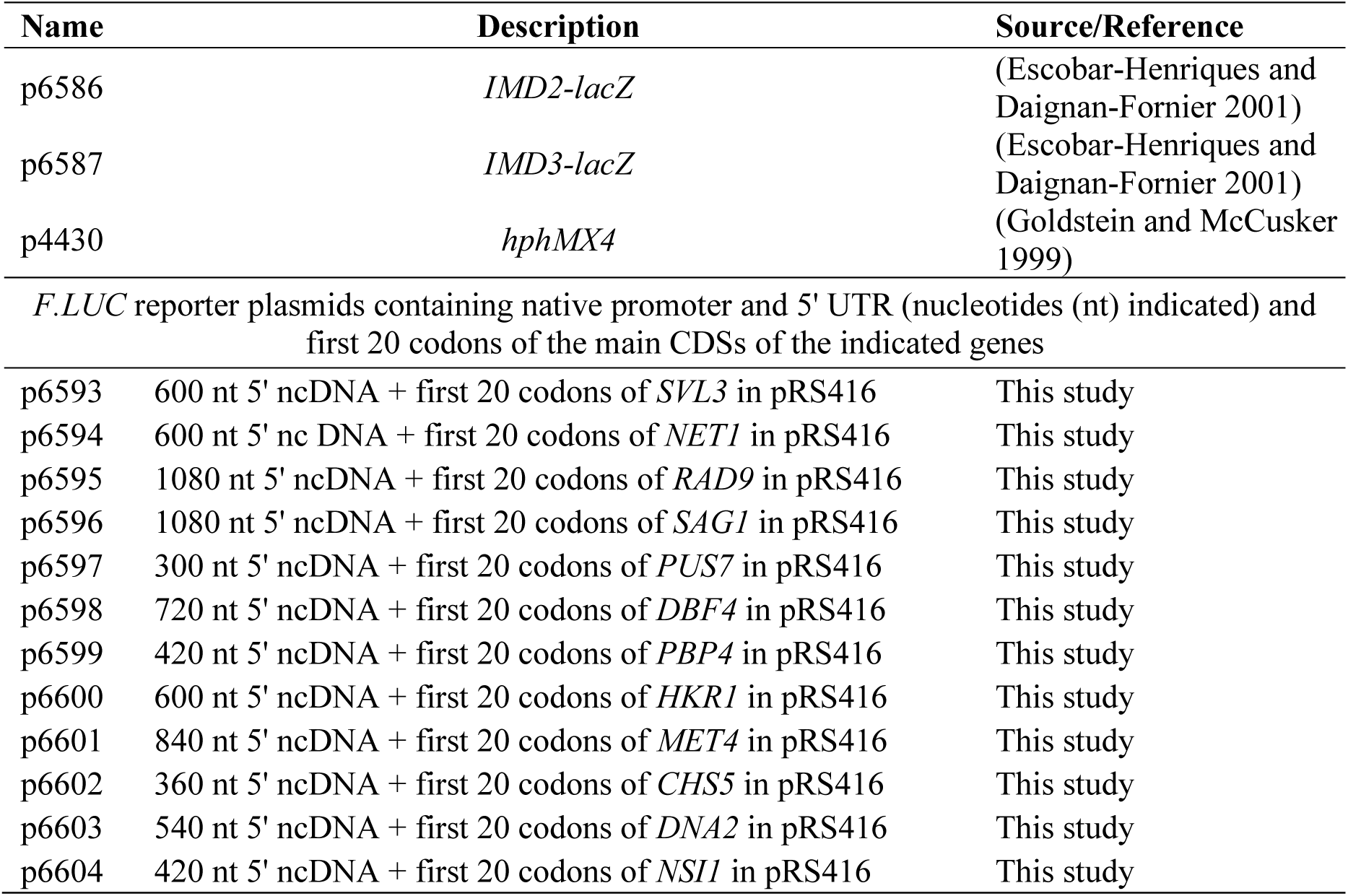
Plasmids used in this study.

**Table 3.**
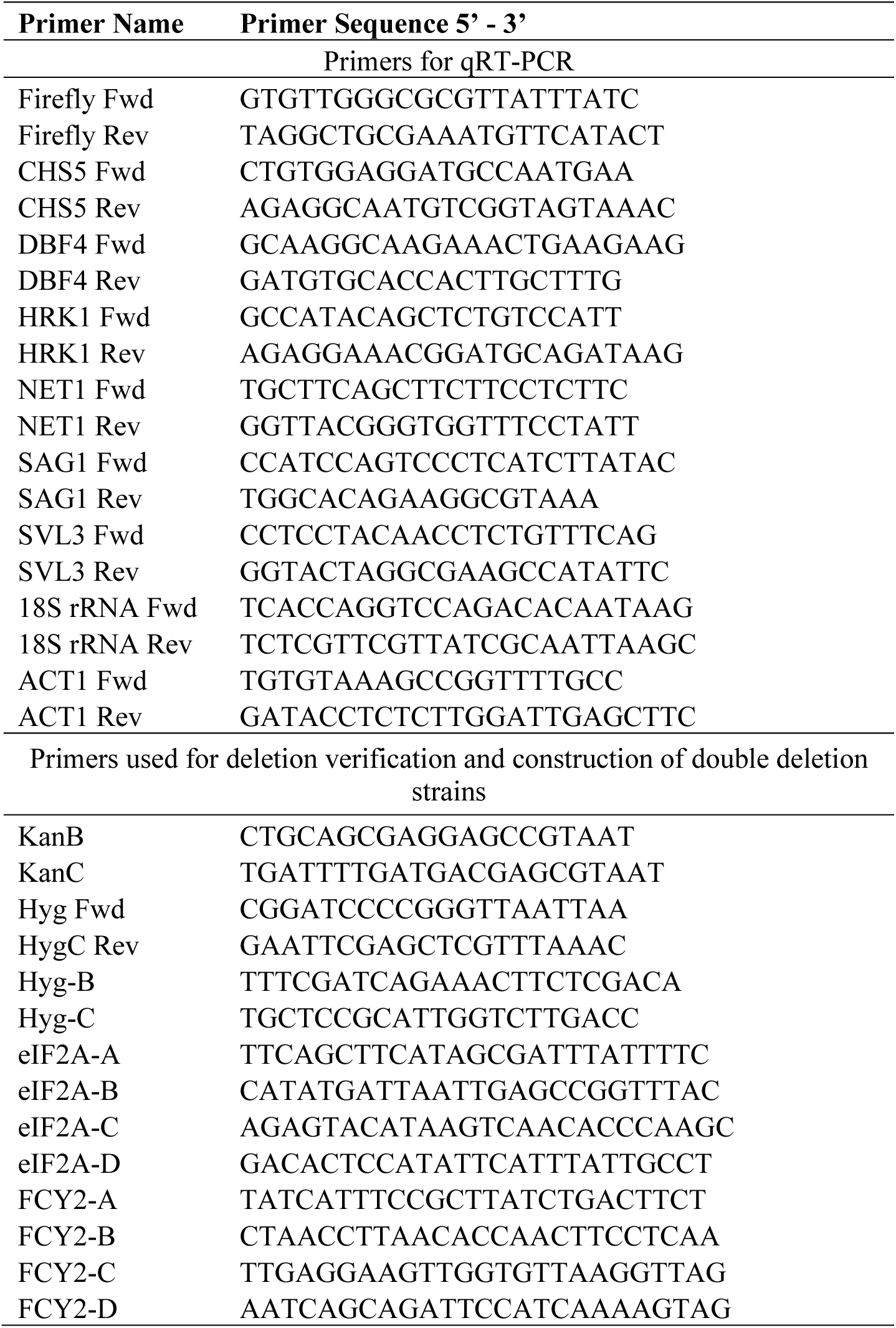
Primers used in this study.

All plasmids employed in this study are listed in Table 2. Plasmids p6593-p6604 were constructed by LifeSct LLC by synthesizing DNA fragments containing the promoter, 5’ UTR, and first 20 codons of the main CDS of the candidate genes and using them to replace the corresponding fragment of p6029 (Sen et al. 2015), fusing the first 20 codons of the candidate genes to the *F.LUC* CDS.

### Preparation of ribosome footprint (RPF) and RNA-seq sequencing libraries

Ribosome profiling and RNA-seq analyses were carried out in parallel on strains BY4742 (WT) and F2247 (*eIF2AΔ*), with two biological replicates for each strain, as described previously (Gaikwad et al. 2021). In brief, both strains were grown at 30°C in SC until reaching log phase at A_600_ = 0.5-0.6 for untreated cells, or to A_600_ = 0.5-0.6 in SC lacking isoleucine and valine (SC-Ile/Val) and treated with SM at 1 µg/mL for 25 min for SM-treated cells. Cells were harvested by high-speed vacuum filtration and snap-frozen in liquid nitrogen. Cell lysis was performed using a freezer mill in the presence of lysis buffer containing 500 μg/mL of cyclohexamide, and RPFs were prepared by digesting cell lysates with RNase I. The 80S monosomes were resolved by sedimentation through a 10-50% sucrose gradient. The RPFs were purified from the monosomes using hot phenol-chloroform extraction. After size selection and dephosphorylation steps, a Universal miRNA cloning linker was attached to the 3’ ends of the footprints. This was followed by reverse transcription, circular ligation, rRNA subtraction, PCR amplification of the cDNA library, and DNA sequencing using an Illumina HiSeq system at the NHLBI DNA Sequencing and Genomics Core at NIH (Bethesda, MD).

For RNA-seq library preparation, total RNA was extracted and purified from aliquots of the same snapped-frozen cells lysates described above using hot phenol-chloroform extraction. Five µg of randomly fragmented total RNA was used for library generation and sequencing, similar to steps mentioned above except that the Ribo-Zero Gold rRNA Removal Kit (Illumina; MRZ11124C) was employed to remove rRNA after linker-ligation.

### Differential gene expression and uORF translation analysis of ribosome profiling data

Processing and analysis of sequence libraries of RPFs or total mRNA fragments, including Wiggle track normalization to the total number of mapped reads for viewing RPF or RNA reads in the IGV browser, were conducted exactly as described previously (Gaikwad et al. 2021). In brief, sequencing reads were trimmed and noncoding RNA sequences were eliminated by aligning trimmed FASTA sequences to the *S. cerevisiae* ribosomal database using Bowtie (Langmead et al. 2009). The remaining reads were mapped to the yeast genome using TopHat (Trapnell et al. 2009), and DESeq2 was used for statistical analysis of mRNA reads, RPFs, and TEs (Love et al. 2014). The R script employed for DESeq2 analysis of TE changes can be found on Github (https://github.com/hzhanghenry/RiboProR; hzhanghenry, 2023; Kim et al., 2019). Wiggle files were visualized using IGV 2.4.14 at (http://software.broadinstitute.org/software/igv/) (Robinson et al. 2011).

Relative ribosome occupancies (RRO) for genes containing translated uORFs were calculated by dividing the RPF counts in the uORF by the RPF counts of the main CDS. DESeq2 was employed to conduct statistical analysis on changes in RRO values between the two biological replicates of different genotypes. uORFs with mean RPF counts below two or mean CDS RPF counts below 32 in the combined samples (two replicates each) were excluded from the analysis. The compilations of annotated translated uORFs (Martin-Marcos et al. 2017), evolutionarily conserved translated uORFs (Spealman et al. 2018), or functional uORFs whose elimination by start codon mutation conferred a consistent and statistically significant alteration in translation of the downstream GFP CDSs in reporter constructs (May et al. 2023), were all published previously and are provided in Figure 6-source data 5-7.

Tripeptide pause scores were computed as described previously (Meydan et al. 2023). Briefly, this involved dividing the 80S reads per million (rpm) of a three-amino acid motif by the average rpm in the surrounding region (±50 nt around each motif). Sites smaller than the ±50 nt window were excluded from the analysis. To generate average pause scores, we calculated the mean of individual pause scores for each tripeptide motif across the translatome. Motifs represented in the genome less than 100 times were excluded to reduce noise in the analysis.

### Yeast biochemical methods

β-galactosidase activities were assayed in whole cell extracts (WCEs) using a modified version of a protocol described previously (Moehle and Hinnebusch 1991) in which the cleavage of ONPG was determined manually after a 30 min incubation with the WCE. Expression of luciferase was assayed in WCEs as previously described (Gaikwad et al. 2021) using the Dual-Luciferase Reporter Assay System (Promega) following the supplier’s protocol, and luciferase activities were normalized to total protein levels in the WCEs extracts determined using the Bradford assay kit (Bio-Rad Laboratories).

### Polysome profile analysis and measurement of TEs of individual mRNAs

Cells were cultured as described in the figure legends, harvested by centrifugation, and WCEs were prepared by vortexing cell pellets, applying eight cycles of vortexing for 30s followed by incubation on ice for 30s, using two volumes of glass beads in ice-cold 1X breaking buffer (20 mM Tris-HCl (pH 7.5), 50 mM KCl, 10 mM MgCl2, 1 mM DTT, 200 μg/mL heparin, 50 μg/mL cycloheximide, and 1 Complete EDTA-free Protease Inhibitor cocktail Tablet (Roche)/50 mL buffer). Fifteen A_260_ units of WCEs were layered on a pre-chilled 10-50% sucrose gradient and centrifuged at 40,000 rpm for 2 h at 4°C in a SW41Ti rotor (Beckman). Gradient fractions were continuously scanned at A_260_ using the BioComp Gradient Station and polysome to monosome (P/M) ratios were calculated using ImageJ software for at least two biological replicates.

To determine polysome association of individual mRNAs and calculate their TEs, 20 A_260_ units of WCEs from each of three replicate cultures were sedimented through a 10-50% sucrose gradient by centrifugation at 35,000 rpm for 2.4 h and the gradient fractions were scanned at A_260_ nm. The fractions containing 80S monosomes or different polysomal species (2-mers, 3-mers etc.) were collected using “the advanced fraction” function of the BioComp fractionator (Figure 3-figure supplement 1). RNA was extracted from 1/5^th^ of the total volume of input WCEs and from 300 µl of each set of pooled fractions containing 80S or different polysomal species using QIAzol lysis Reagent (Qiagen) according to the manufacturer’s protocol. Reverse transcription (RT) was done using SuperScript III First-Strand Synthesis SuperMix (Invitrogen) with random hexamers, using five μg of RNA for each reverse transcription (RT) reaction. Transcript levels were quantified by qPCR using the Brilliant III Ultra-Fast SYBR Green qPCR Master Mix (Agilent Technologies) on a Roche LightCycler 96 Instrument. Primers listed in Table S3 were used to measure the individual mRNAs and 18S rRNA. The absolute mRNA/18S rRNA levels were calculated using the 2^-CT^ method and corrected to account for differences in the proportions of the pooled fractions from which RNA was isolated by multiplying by a factor calculated by dividing the total fraction volume (in µl) by 300 µl. They were further corrected to account for differences in the proportion of the total RNA employed for RT by multiplying by a factor calculated as 25 µl (the total volume of extracted mRNA) divided by the volume of the pooled fraction (in µl) containing 5 µg of RNA. To account for potential losses during RNA extraction, RNA recovery normalization factors were calculated for each fraction by determining the proportion of total A_260_ units across all gradient fractions that is present in each set of pooled fraction (calculated from the A_260_ trace obtained during gradient fractionation) and dividing the results by the proportion of total 18S rRNA across the gradient found in the corresponding pooled fraction. A second normalization factor was calculated to correct for losses in recovery of monosomes/polysomes in the gradient separations of fixed amounts of input WCEs by calculating the total A_260_ units found in monosomes/polysomes for each gradient, determining the mean value for all of the gradients/samples analyzed in parallel, and dividing the mean value by the value determined for each gradient. (This correction assumes that WT and *eIF2AΔ* cells have the same total amounts of monosomes/polysomes per A_260_ of WCE, as indicated by our repeated polysome profiling of biological replicates depicted in Figure 1A.) The absolute amounts of mRNAs measured in each pooled fraction were multiplied by both normalization factors to obtain the normalized mRNA levels for each pooled fraction, which was multiplied by the number of ribosomes per mRNA in that pooled fraction (i.e., 1 for monosomes, 2 for 2-mer polysomes, etc.) and summed across the gradient fractions to calculate the total number of ribosomes translating the mRNA. To calculate TEs, this last quantity was divided by amount of input mRNA measured in the starting WCE normalized to level of *ACT1* mRNA. For each of three biological replicate cultures of each strain, WT or *eIF2AΔ* mutant, we determined the ratio of TE in the presence versus absence of SM and calculated the mean and SEM values for ΔTEs.

### Statistical analyses and data visualization

Notched box plots were created using a web-based tool (http://shiny.chemgrid.org/boxplotr/). Scatterplots and volcano plots were generated using the scatterplot function in Microsoft Excel. Hierarchical cluster analysis of TE changes in mutants/conditions was performed using the R heatmap.2 function from the ’gplots’ library using the default hclust hierarchical clustering algorithm. Smoothed scatterplots were computed and plotted using the ggplot2 package in R. Calculation of Spearman’s correlation coefficients and student’s t-tests were performed using built-in features of Microsoft Excel. The Mann-Whitney U test and p-values for Pearson’s correlation were computed using the R Stats package in R.

## Data availability

Sequencing data from this study have been deposited to the NCBI Gene Expression Omnibus (GEO) under GEO accession number GSE241473.

## ACKNOWLEDGEMENTS

We thank Sezen Meydan for performing the tripeptide pause score analysis and Bertrand Daignan-Fornier for gifts of reporter plasmids. We are grateful to members of our laboratory and those of the Lorsch, Dever, and Guydosh labs for many helpful suggestions. This work was supported by the Intramural Research Program of the National Institutes of Health.

## SUPPLEMENTARY FIGURE LEGENDS

**Figure 1-figure supplement 1.**
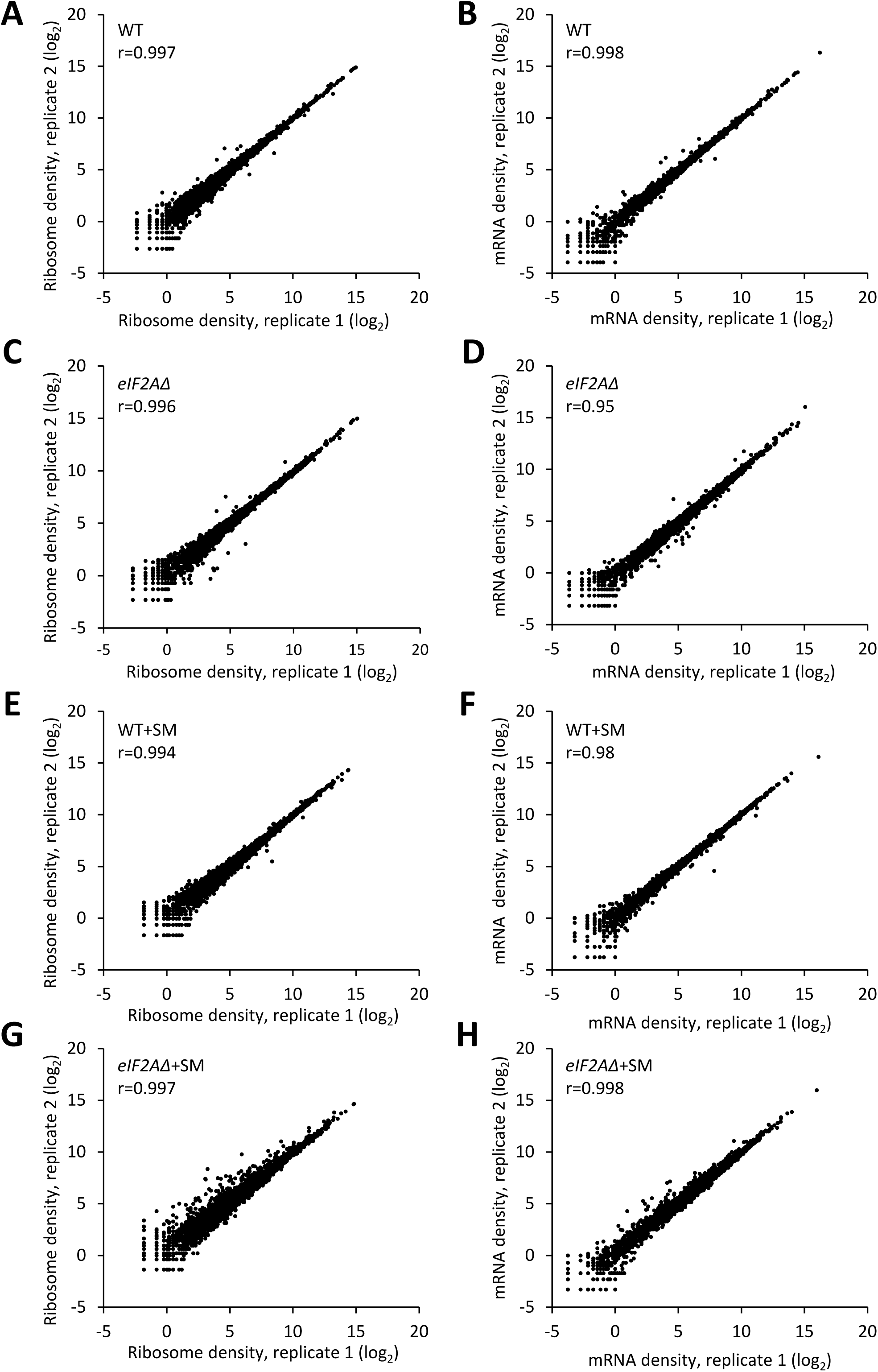
High reproducibility between biological replicates of ribosome footprint profiling and RNA-seq analyses. **(A-H)** Scatterplots depict the RPF (A, C, E, G) or mRNA (B, D, F, H) read densities for all expressed mRNAs across biological replicates of the WT (A, B), the *eIF2AΔ* mutant (C, D) SM-treated WT (E, F) and SM-treated *eIF2AΔ* mutant (G, H). The read densities were calculated by mapping the reads to the CDS of each gene and expressed as reads per million mapped reads (RPM) in individual libraries of biological replicates. The Pearson’s coefficient (r) is indicated in each plot, quantifying the degree of correlation between the replicate datasets.

**Figure 2-figure supplement 1.**
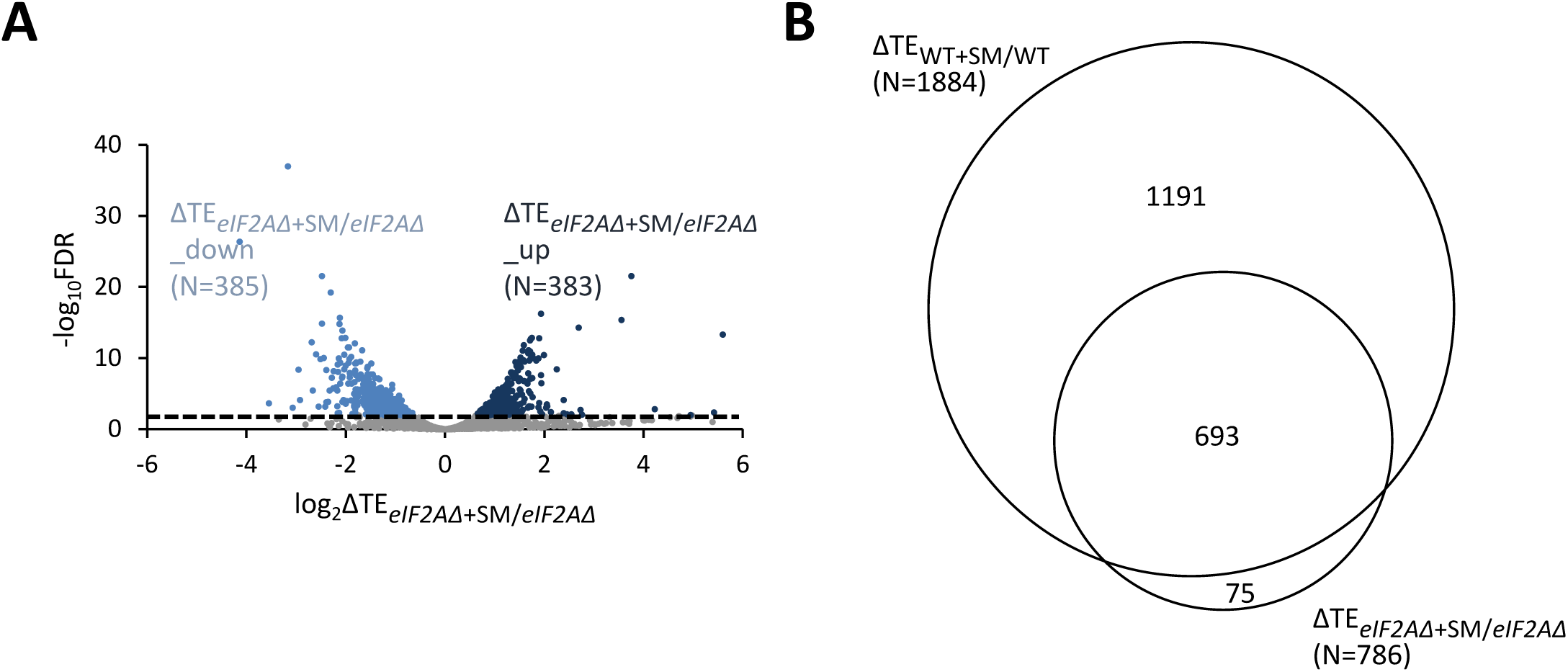
Relative TE changes evoked by increased eIF2α phosphorylation in cells lacking eIF2A are broadly similar to relative TE changes conferred by increased eIF2α phosphorylation in WT cells. **(A)** Volcano plot as in Figure 2A showing the log_2_ ratios of TEs in SM-treated *eIF2AΔ* versus untreated *eIF2AΔ* cells (ΔTE*_eIF2AΔ_*_+SM/*eIF2AΔ*_ values) for the 5426 mRNAs with evidence of translation. The dotted line marks the 1% FDR threshold. Genes showing a significant increase (ΔTE*_eIF2AΔ_*_+SM/*eIF2AΔ*__up) or decrease (ΔTE*_eIF2AΔ_*_+SM/*eIF2AΔ*__down) in TE in SM-treated *eIF2AΔ* versus *eIF2AΔ* mutant cells at FDR < 0.05, are plotted in dark and light blue circles, respectively. **(B)** Proportional Venn diagram showing overlap between the 1884 mRNAs identified in Figure 2B and the 786 mRNAs identified in Figure 2-figure supplement 1A.

**Figure 3-figure supplement 1.**
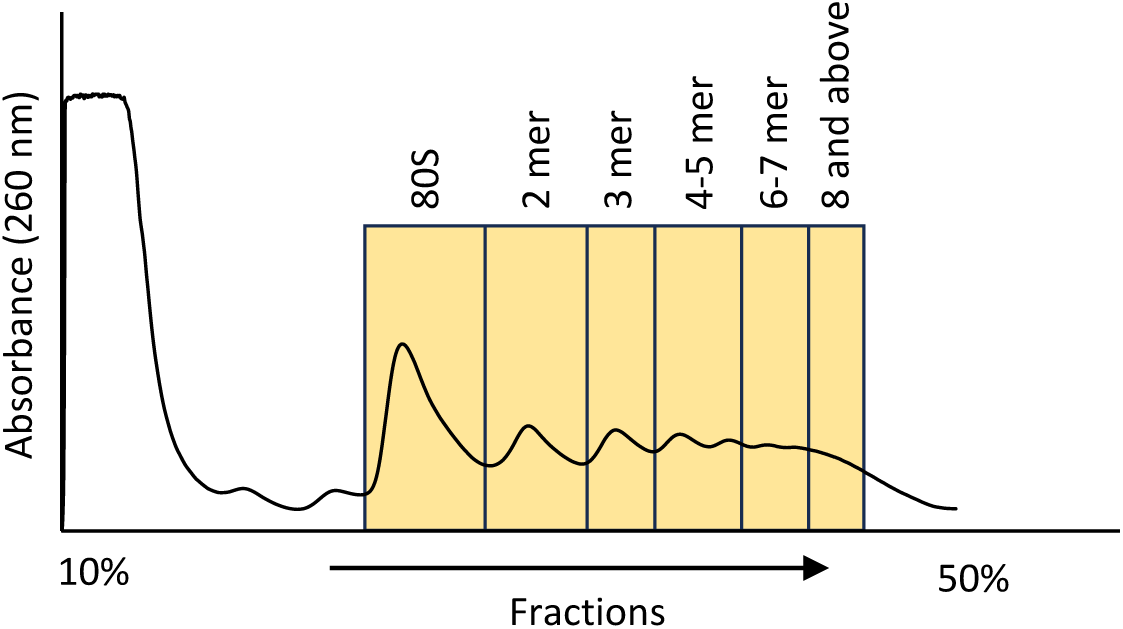
Representative separation of polysomes by sedimentation through a sucrose density gradient in the experiment depicted in. Figure 3E. The A_260_ values were determined continuously during fractionation of the gradient. Fractions pooled for isolation of RNA from 80S monosomes or the various polysomal species are indicated by boxes.

**Figure 6-figure supplement 1.**
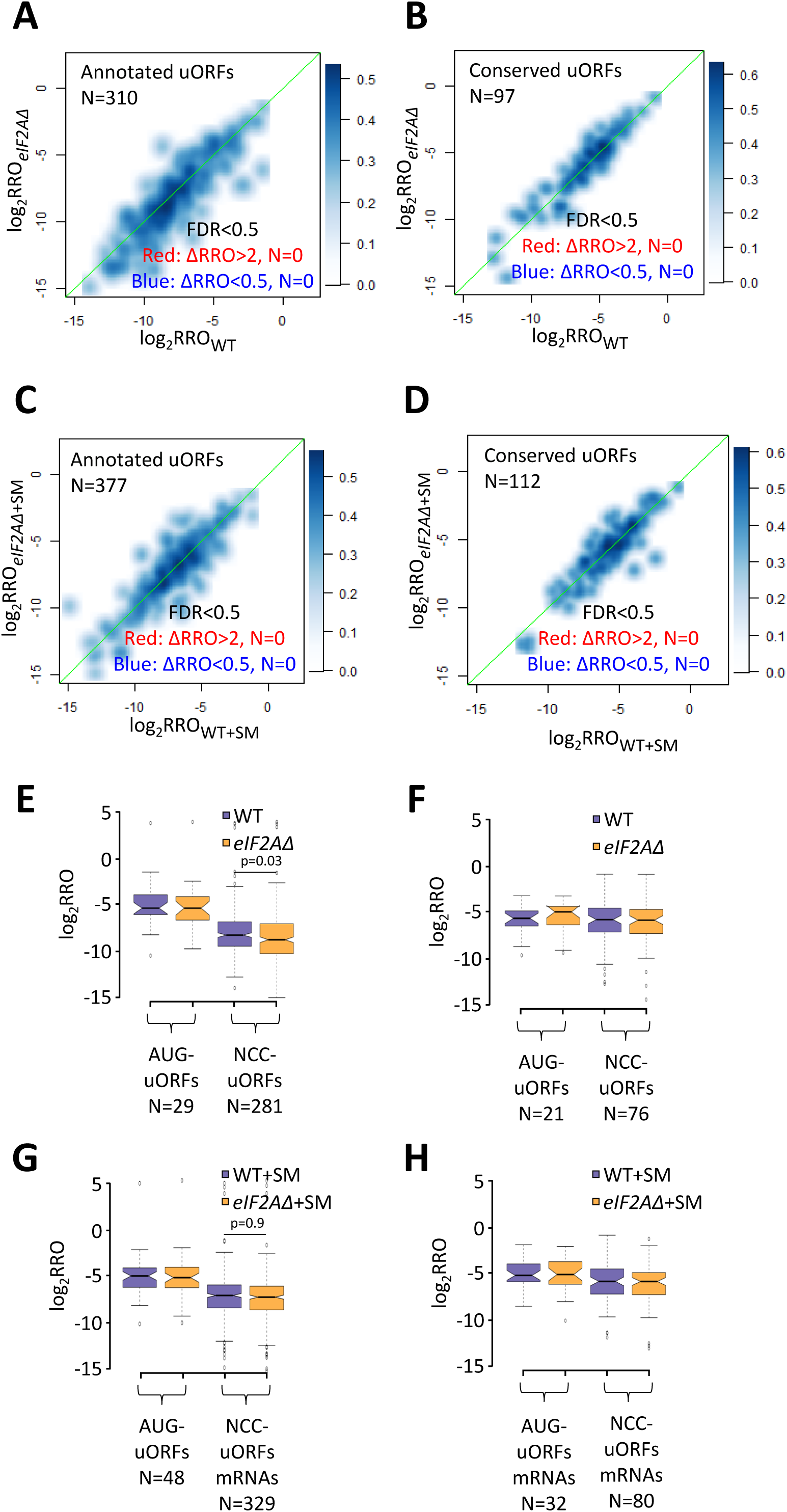
eIF2A plays a minimal in regulating uORF-mediated translation. **(A-B)** Smoothed scatterplots displaying the relationship between log_2_RRO_WT_ (x-axis) and log_2_RRO*_eIF2AΔ_* (y-axis) for all mRNAs containing annotated AUG- or NCC-uORFs (A) or evolutionarily conserved AUG- or NCC-uORFs (B) in WT versus *eIF2AΔ* cells without SM treatment. No mRNAs showed ≥2-fold changes in RRO in the *eIF2AΔ* mutant versus WT cells at FDR < 0.5. **(C-D)** Smoothed scatterplots displaying the relationship between log_2_RRO_WT+SM_ (x-axis) versus log_2_RRO*_eIF2AΔ_*_+SM_ (y-axis) for the same mRNAs analyzed in (A)-(B) but in the presence of SM. Again, no mRNAs showed ≥2-fold changes in RRO in the *eIF2AΔ* mutant versus WT cells at FDR < 0.5. **(E-H)** Notched box plot displaying log_2_RRO values for all mRNAs containing annotated AUG- or NCC-uORFs (E, G) or evolutionarily conserved AUG- or NCC-uORFs (F, H) in untreated WT and *eIF2AΔ* mutant (E-F) or SM-treated WT and *eIF2AΔ* mutant (G-H). The y-axis scale was expanded by omitting a few outliers. Statistical significance determined using the Mann-Whitney U test is shown for the bracketed comparisons in panels E & G.

**Figure 6-figure supplement 2.**
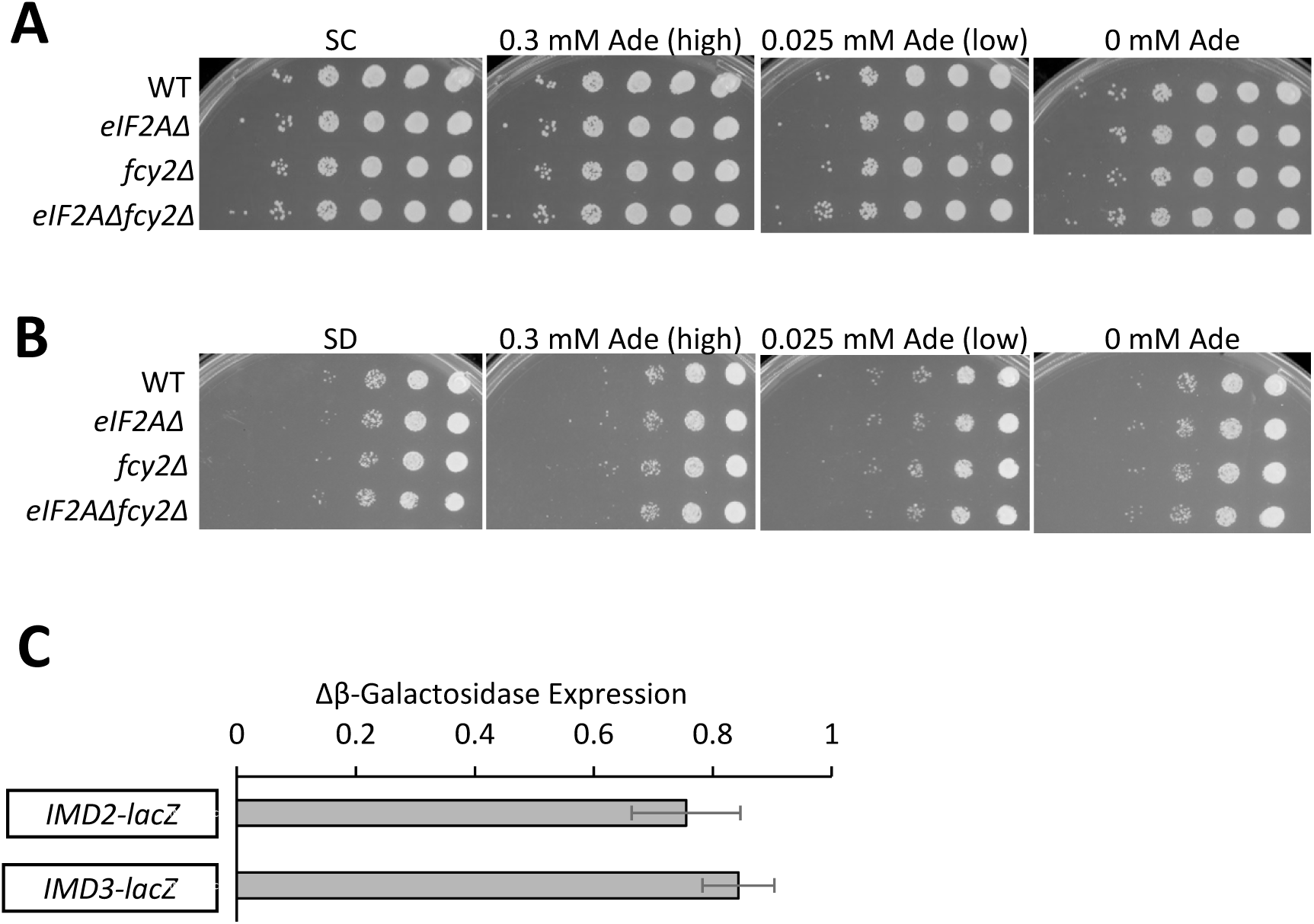
Lack of genetic interaction between eIF2A and the purine salvage pathway. **(A-B)** Cell spotting assays were performed on SC plates (A) or SD (B) plates to assess the growth of WT, *eIF2AΔ*, *fcy2Δ* and *eIF2AD fcy2D* strains. Ten-fold serial dilutions of saturated cultures were applied to SC or SD plates supplemented with the indicated concentrations of adenine and incubated at 30°C for 2 days. **(C)** WT and *eIF2AΔ* strains were transformed with the indicated *lacZ* reporter plasmids-IMD2 and IMD3. The transformants bearing the reporter plasmids-IMD2 containing IMD2 coding sequences along with 1186 bp upstream region while IMD3 contains IMD3 coding sequences containing 555 bp upstream region-were grown in SC-Ura to saturation. The cultures were then diluted in fresh SC-Ura containing 0.015 mM concentration of Adenine and grown for 6 h to A_600_ of ∼1.0. WCEs were prepared and assayed for β-galactosidase activities in units of nmol of ONPG cleaved per mg of protein per min. The results represent the fold change of means and ±SEMs of activities calculated from three independent transformants.

## FIGURE SOURCE DATA

**Figure 2-source data 1.**

Spreadsheet tabulates the log_2_ ratios of TEs in *eIF2AΔ* versus WT cells (log_2_ΔTE*_eIF2AΔ_*_/WT_ values) for each mRNA and the corresponding FDR determined by DESeq2 analysis of ribosome profiling and parallel RNA Seq data for the 5340 mRNAs with evidence of translation (Figure 2A).

**Figure 2-source data 2.**

Spreadsheet tabulates the log_2_ ratios of TEs in WT+SM cells versus WT cells (log_2_ΔTE_WT+SM/WT_ values) and the corresponding FDR determined by DESeq2 analysis of ribosome profiling and parallel RNA Seq data for the 5441 mRNAs with evidence of translation (Figure 2B & Figure 2-figure supplement 1B).

**Figure 2-source data 3.**

Spreadsheet tabulates the log_2_ ratios log_2_ΔTE values for the indicated mutant/condition for the indicated

mRNA groups identified in Figure 2B (Figure 2D).

**Figure 2-figure supplement 1-source data 1.**

Spreadsheet tabulates the log_2_ ratios of TEs in SM-treated *eIF2AΔ* versus untreated *eIF2AΔ* cells (ΔTE*_eIF2AΔ_*_+SM/*eIF2AΔ*_ values) for each mRNA and the corresponding FDR determined by DESeq2 analysis of ribosome profiling and parallel RNA Seq data for the 5426 mRNAs with evidence of translation (Figure 2-figure supplement 1).

**Figure 3-source data 1.**

Spreadsheet tabulates the log_2_ ratios of TEs in *eIF2AΔ* cells treated with SM versus WT cells treated with SM (log_2_ΔTE*_eIF2AΔ_*_+SM/WT+SM_ values) for each mRNA and the corresponding FDR determined by DESeq2 analysis of ribosome profiling and parallel RNA Seq data for the 5482 mRNAs with evidence of translation (Figure 3A).

**Figure 3-source data 2.**

Spreadsheet tabulates the log_2_ΔTE values for the indicated mutant/condition for the 32 mRNAs in the group ΔTE*_eIF2AΔ_*_+SM/WT+SM_down_ defined in Figure 3A (Figure 3B).

**Figure 3-source data 3.**

Spreadsheet tabulates the log_2_ΔTE values for the indicated mutant/condition for the 32 mRNAs in the group ΔTE*_eIF2AΔ_*_+SM/WT+SM_down_ defined in Figure 3A (Figure 3C).

**Figure 3-source data 4.**

Spreadsheet tabulates the changes in luciferase activity expressed from F.LUC reporters calculated for WT+SM versus WT and *eIF2AΔ+SM* versus *eIF2AΔ* for three biological replicates (Figure 3D).

**Figure 3-source data 5.**

Spreadsheet 1, “raw CT values for mRNA from qRT”, tabulates the raw CT values for mRNA from qRT reactions for three biological replicates-a, b, and c. Spreadsheet 2, “Dilution factor exemplar WT_a”, provides an example file for one of the biological replicate WT (wild type) samples, illustrating how to calculate the dilution factor. Spreadsheet 3, “% 18S rRNA”, provides an example file illustrating how to calculate the dilution factor. Spreadsheet 4, “Area % & Polysome norm factor”, tabulates following parameters for all the biological replicates: Area under peaks from UV trace; SUM 80S+Polysomes; Average SUM 80S+Polysomes; Polysome recovery normalisation factor; and % total Area under 80S + Polysomes from UV trace in each peak. Spreadsheet 5, “rRNA normalisation factor”, illustrates the calculations required to determine the rRNA normalization factor. Spreadsheet 6, “SAG1”, illustrates the calculations required to determine ΔMonosome-Polysome/Total RNA (See materials and methods section for details) (Figure 3E).

**Figure 6-source data 1.**

Spreadsheet 1-3, “Fig 6A(i-iii)”, tabulates the lists and log_2_ΔTE values for the indicated mutant/condition for the annotated AUG- or NCC-uORFs (Fig 6Ai), conserved AUG- or NCC-uORFs (Fig 6Aii), or single functional inhibitory AUG-uORFs (Fig 6Aiii) (Figure 6A).

**Figure 6-source data 2.**

Spreadsheet 1-3, “Fig 6B(i-iii)”, tabulates the lists and log_2_ΔTE values as in (Figure 6-source data 1) for the subsets of the same mRNA groups analyzed there exhibiting > 1.41-fold increases in TE in SM-treated versus untreated WT cells (Figure 6B).

**Figure 6-source data 3.**

Spreadsheet 1-3, “Fig 6C(i-iii)”, tabulates the lists and log_2_ΔTE values as in (Figure 6-source data 1) for the subsets of the same mRNA groups analyzed there exhibiting > 1.41-fold decreases in TE in SM-treated *eIF2AΔ* versus SM-treated WT cells (Figure 6C).

**Figure 6-source data 4.**

Spreadsheet tabulates lists of the 514 mRNAs bearing functional AUG or NCC-uORFs, 17 mRNAs identified in Figure 3A showing evidence for a conditional requirement for eIF2A when eIF2 function is reduced by SM i.e., ΔTE*_eIF2AΔ_*_+SM/WT+SM__down* (N=17) group, and overlap of mRNAs with functional AUG- or NCC-uORFs (N=514) and ΔTE*_eIF2AΔ_*_+SM/WT+SM__down (N=17) (Figure 6D).

**Figure 6-source data 5.**

Spreadsheet tabulates the list of annotated AUG- or NCC-uORFs, chromosome coordinates, start codon of uORF, distances of the uORF AUG from the 5’ end of the mRNA and the main CDS start codon, and the gene name.

**Figure 6-source data 6.**

Spreadsheet tabulates the list of conserved AUG- or NCC-uORFs, chromosome coordinates, start codon of uORF, distances of the uORF AUG from the 5’ end of the mRNA and the main CDS start codon, and the gene name.

**Figure 6-source data 7.**

Spreadsheet tabulates the functional uORFs , chromosome coordinates, start codon of uORF, distances of the uORF AUG from the 5’ end of the mRNA and the main CDS start codon, and the gene name (May et al. 2023).

**Figure 6-figure supplement 1-source data 1.**

Spreadsheet 1 tabulates the log_2_ ratios of following parameters for all the expressed annotated uAUG or NCC-uORFs listed in col. A in *eIF2AΔ* versus WT cells: Relative Ribosome Occupancy (RRO) in *eIF2AΔ* versus WT cells (RRO Change), RRO for WT, and RRO of *eIF2AΔ* (Figure 6-figure supplement 1A & E). Figure 6-figure supplement 1-source data 2.

Spreadsheet 1 tabulates the log_2_ ratios of following parameters for all the evolutionarily conserved expressed uAUG or NCC-uORFs listed in col. A in *eIF2AΔ* versus WT cells: Relative Ribosome Occupancy (RRO) in *eIF2AΔ* versus WT cells (RRO Change), RRO for WT, and RRO of *eIF2AΔ* (Figure 6-figure supplement 1B & F).

**Figure 6-figure supplement 1-source data 3.**

Spreadsheet 1 tabulates the log_2_ ratios of following parameters for all the expressed annotated uAUG or NCC-uORFs listed in col. A in SM-treated WT and *eIF2AΔ* mutant: Relative Ribosome Occupancy (RRO) in *eIF2AΔ* treated with SM versus WT cells treated with SM (RRO Change), RRO for WT treated with SM, and RRO of *eIF2AΔ* treated with SM (Figure 6-figure supplement 1C & G).

**Figure 6-figure supplement 1-source data 4.**

Spreadsheet 1 tabulates the log_2_ ratios of following parameters for all the evolutionarily conserved expressed uAUG or NCC-uORFs listed in col. A in *eIF2AΔ* treated with SM versus WT cells treated with SM: Relative Ribosome Occupancy (RRO) in SM-treated WT and *eIF2AΔ* mutant (RRO Change), RRO for WT treated with SM, and RRO of *eIF2AΔ* treated with SM (Figure 6-figure supplement 1D & H).

**Figure 6-figure supplement 1-source data 5.**

Spreadsheet 1 tabulates the β-galactosidase activities in units of nmol of ONPG cleaved per mg of protein per min in WT and *eIF2AΔ* strains. Additionally, the spreadsheet provides the fold change of means and the associated ±SEMs (Standard Error of the Means) of these activities. These values have been calculated based on data obtained from three independent transformants for each condition (Figure 6-figure supplement 2B).

